# KELPE: knock-in exchangeable dual landing pad embryonic stem cells enable efficient screening of synthetic gene circuits

**DOI:** 10.64898/2026.03.22.713470

**Authors:** Aisling Fairweather, Yana Slavova, Mattias Malaguti

**Affiliations:** Institute for Regeneration and Repair, College of Medicine and Veterinary Medicine, University of Edinburgh, Edinburgh EH16 4UU, UK; Centre for Engineering Biology, Institute of Quantitative Biology, Biochemistry and Biotechnology, School of Biological Sciences, College of Science and Engineering, University of Edinburgh, Edinburgh EH9 3FF, UK

**Keywords:** Pluripotent stem cells, synthetic biology, transgenesis, genetic circuits, synNotch

## Abstract

The establishment of genetic circuits in pluripotent stem cells (PSCs) allows to model and manipulate developmental events. However, prototyping complex circuitry remains challenging, due to limitations in screening circuit components and transgene silencing. Here, we introduce KELPE: PSCs with two silencing-resistant insulated genomic landing pads targeted to genomic safe harbour sites.

KELPE cells enable the stable integration of multiple transgenes into the same genomic region, facilitating fair comparisons of genetic circuit components. We demonstrate this by fine-tuning “synthetic neighbour-labelling” technologies. We first generate optimised PUFFFIN PSCs, which report on cell-cell interactions by fluorescently labelling wild-type neighbours. We then generate new synNotch “receiver” PSCs, which can trigger expression of any transgene following interaction with a synthetic ligand presented by “sender” cells of interest. We describe an optimised circuit syntax that abolishes ligand-independent transgene induction in receiver PSCs, and showcase this by synthetically programming cell death in receiver cells engineered to express a toxin following interaction with sender cells.

In summary, we describe a new cell line that facilitates silencing-resistant transgene expression and prototyping of synthetic biology tools in a developmentally-relevant model.

## INTRODUCTION

Synthetic developmental biology is a discipline that seeks to recapitulate and control developmental events by employing synthetic biology tools and engineering principles (Davies, 2017; McNamara et al., 2023; Schlissel and Li, 2020). It is heavily reliant on the establishment of complex genetic circuits consisting of multiple transgenic cassettes (Ebrahimkhani and Ebisuya, 2019; Ho and Morsut, 2021; Santorelli et al., 2019). Pluripotent stem cells (PSCs) are amenable to transgenesis, and are therefore a popular developmental model for the establishment of genetic circuits (Lienert et al., 2013; Prochazka et al., 2022; Saxena et al., 2016). However, this process is accompanied by a series of technical challenges.

When developing genetic circuits, a key step is represented by the comparison of different genetic elements, such as promoters, terminators or expression cassettes. This can be achieved through several approaches, a common one being the transient delivery of genetic circuits on non-integrating plasmids, which allows for rapid prototyping of circuit behaviour (De Carluccio et al., 2024; Scheller et al., 2018; Strittmatter et al., 2022; Zhou et al., 2023). However, this strategy relies on comparable transfection efficiency both between different conditions and in cells within the same culture vessel, which is usually unattainable (Bell et al., 2007; Di Blasi et al., 2021; Vitor et al., 2018).

An alternative approach is to stably integrate different circuit components into random regions within the genome of host cells; this allows researchers to compare circuit behaviour efficiency across polyclonal populations or across monoclonal cell lines (Malaguti et al., 2022; McNamara et al., 2024; Soliman et al., 2025). However, expression of transgenes integrated into different regions of the genome is affected by adjacent endogenous DNA elements: transgene expression can be boosted by nearby enhancer elements, or dampened by repressive epigenetic marks (Cranston et al., 2001; Dobie et al., 1996; Gierman et al., 2007; Williams and Wagner, 2000). This confounds the comparison of different components of genetic circuits integrated into different genomic regions.

A third approach is the integration of genetic circuits into “safe harbour” genomic sites, such as the mouse *Gt(ROSA)26Sor* (“*Rosa26*”) locus (Friedrich and Soriano, 1991). These safe harbour loci allow ubiquitous transgene expression across tissues and developmental stages (Kisseberth et al., 1999; Srinivas et al., 2001; Zambrowicz et al., 1997), and therefore provide ideal comparison platforms for different genetic circuit components. However, targeting DNA constructs to specific genomic sites requires the addition of homology arms to transgenic vectors, increasing their size and consequently decreasing their transfection efficiency (Kreiss et al., 1999; Lesueur et al., 2016; Ribeiro et al., 2012). Furthermore, transgene integration at the correct genomic location needs to be validated for each clonal line, considerably increasing screening time.

A solution to this issue is represented by “genomic landing pads”, DNA constructs that contain stuffer DNA flanked by short specific sites that are recognised by site-specific recombinase enzymes (SSRs). SSRs can catalyse the replacement of the stuffer DNA with any “donor” sequence flanked by the relevant SSR sites, via a process termed “recombination-mediated cassette exchange” (RMCE) (Fong and Ceroni, 2025; Merrick et al., 2018; Turan et al., 2013). This highly efficient process allows for a fair comparison of the performance of different genetic circuit components introduced into the same genomic region of sister cell lines (Matreyek et al., 2017; Matreyek et al., 2020; Zhu et al., 2023). Donor DNA vectors do not require components already integrated into the genome (e.g. homology arms, promoters, terminators, etc.), reducing construct size, and therefore streamlining vector construction and delivery (Gaidukov et al., 2018; Roelle et al., 2023; Shah et al., 2015).

Several research groups have sought to facilitate mouse PSC transgenesis by developing cell lines containing genomic landing pads targeted to safe harbour sites (Chen et al., 2011a; Fitzgerald et al., 2020; Haenebalcke et al., 2013; Hitz et al., 2007; Iacovino et al., 2011; Malaguti et al., 2022; Masui et al., 2005; Rath et al., 2025; Sandhu et al., 2010; Tchorz et al., 2012; Tosti et al., 2018; Tsanov et al., 2012; Zhou et al., 2013). Whilst these lines should in principle allow for rapid comparisons of different genetic circuit components in mouse PSCs, the expression of transgenes integrated into safe harbour sites has been reported to vary considerably across different mouse tissues (Chen et al., 2011b; Yu et al., 2024), suggesting the presence of nearby tissue-specific enhancer elements which may disrupt the desired behaviour of genetic circuits following pluripotent cell differentiation. To address this issue, DNA constructs for random or targeted integration can be flanked by insulator sequences (Ordovás et al., 2015; Rival-Gervier et al., 2013). These are DNA elements that can insulate transgenes from the activity of nearby promoters and enhancers, as well as inhibit transgene silencing by blocking the spread of heterochromatin (Bell et al., 2001; Chung et al., 1993).

A further issue with cell lines carrying a safe harbour locus-targeted genomic landing pad is that it can prove troublesome to deliver complex synthetic circuits to a single genomic location, due to the large size of multi-component vectors affecting efficiency of construct delivery (Kreiss et al., 1999; Lesueur et al., 2016; Ribeiro et al., 2012). To address this issue in human models of development, human PSCs harbouring two identical (Blanch-Asensio et al., 2022; Rosenstein et al., 2024) or two different (Blanch-Asensio et al., 2025) genomic landing pads have recently been described. These tools offer high flexibility in synthetic circuit design and development in a human PSC context.

In this study, we sought to generate a flexible tool to facilitate engineering of synthetic gene circuits in mouse PSCs. Here, we introduce KELPE (Knock-in Exchangeable Landing Pad Embryonic stem cells): the first clonal mouse embryonic stem cell (mESC) line containing two insulated modular genomic landing pads targeted to the two alleles of the safe harbour *Rosa26* locus. Each landing pad encodes a fluorescent transgene that can be replaced by donor DNA sequences flanked by any one of 4 pairs of orthogonal SSR sites, allowing for convenient screening of successful RMCE events (Fig. 1A-C). We demonstrate that transgenes in the genomic landing pads are resistant to silencing, and describe a straightforward method to generate complex vectors for RMCE into both landing pads. We showcase KELPE cells by generating new clonal cell lines harbouring “synthetic neighbour-labelling” circuits, which allow users to fluorescently label the neighbours of any cell of interest (reviewed in Malaguti et al., 2024). We demonstrate how circuit optimisation can lead to optimisation of neighbour labelling by ablating leaky expression of synthetic Notch (synNotch)-inducible transgenes, allowing us to program cell death in response to interaction with a cell of interest.

**Figure 1.**
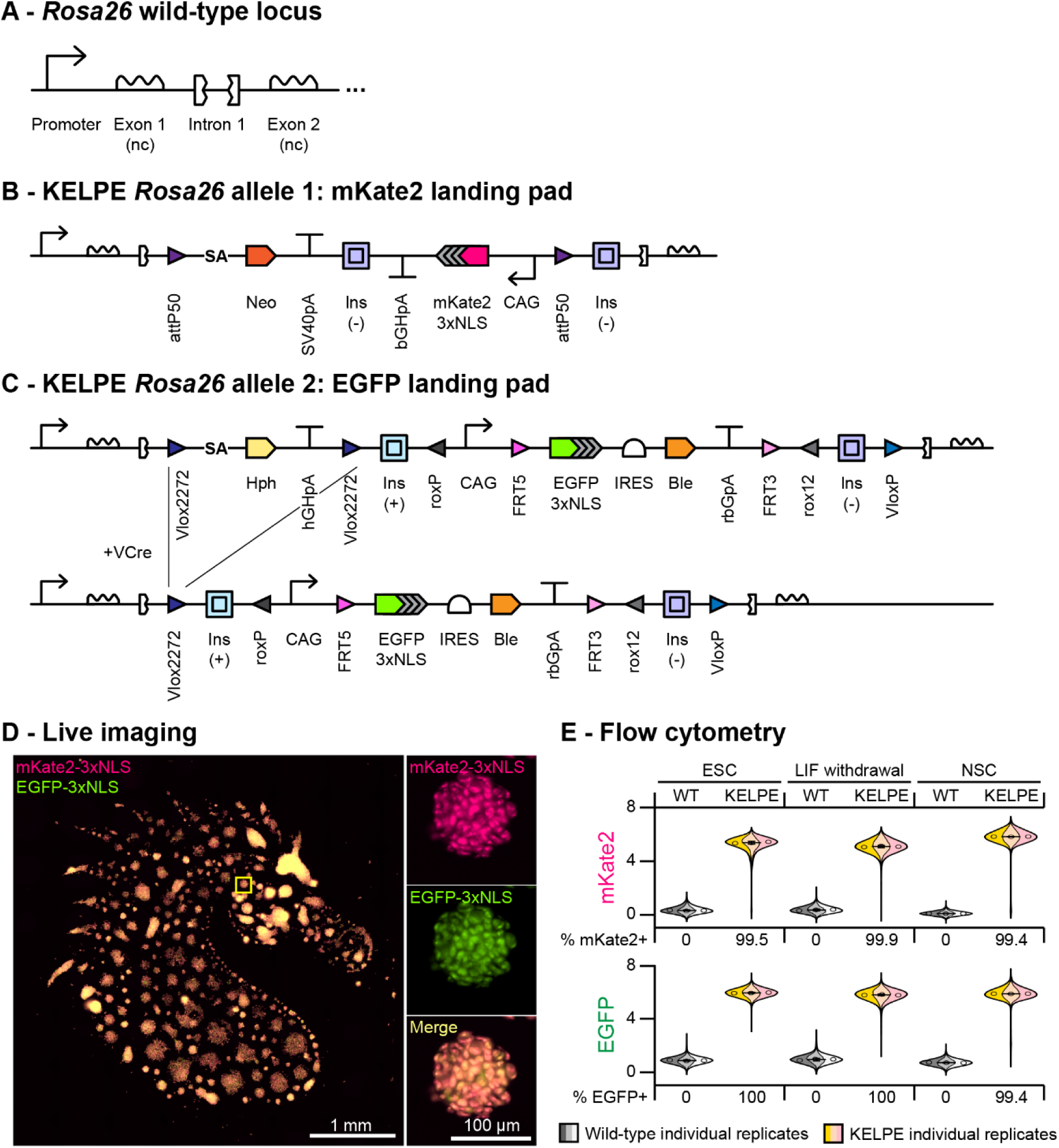
Development and characterisation of knock-in exchangeable landing pad mouse embryonic stem cells (KELPE) (A) Diagram of the wild-type Rosa26 locus. nc: non-coding.sarala(B) Diagram of the Rosa26 locus targeted with the “mKate2” genomic landing pad. SA: splice acceptor; Neo: neomycin phosphotransferase II; SV40pA: SV40 polyadenylation signal sequence; Ins: cHS4 insulator; bGHpA: bovine growth hormone polyadenylation signal sequence.saralaC) Diagram of the Rosa26 locus targeted with the “EGFP” genomic landing pad. Hph: hygromycin phosphotransferase; hGHpA: human growth hormone polyadenylation signal sequence; IRES: internal ribosome entry site; Ble: bleomycin resistance gene; rbGpA: rabbit beta-globin polyadenylation signal sequence.sarala(D) Live imaging of mKate2-3xNLS and EGFP-3xNLS expression in KELPE cells micropatterned in the shape of a kelpie. The panels on the right display a magnified view of the area boxed in yellow in the panel on the left.sarala(E) Flow cytometry analysis of mKate2 and EGFP levels in KELPE cells cultured for a minimum of 28 days in the absence of selective antibiotics in pluripotency culture conditions (“ESC”), in undirected differentiation medium (“LIF withdrawal”) or following differentiation to neural stem cells (“NSC”). Violin Superplots display three individual replicates in different shading within the same distribution. Central line: median; error bars: 95% confidence interval. Ten-thousand cells were analysed tor each replicate.

## RESULTS

### Design and generation of dual landing pad PSCs

We set out to design modular landing pads to target both alleles of the *Rosa26* safe harbour locus in mESCs. This locus consists of a ubiquitous promoter driving expression of a long non-coding RNA (Zambrowicz et al., 1997) (Fig. 1A). Transgenic cassettes can be efficiently targeted into its first intron without the need for nuclease-induced double-strand breaks (Hohenstein et al., 2008; Mort et al., 2014; Nyabi et al., 2009), reducing the risk of mutation or chromosomal rearrangements at off-target genomic sites. Furthermore, transgene insertion at this locus causes no overt phenotype, with targeted mice able to breed to homozygosity (Favero et al., 2018; Platt et al., 2014; Soriano, 1999).

We elected to use degenerate pairs of orthogonal SSR sites which would be recognised by different recombinases (Table S1). We avoided loxP sites (Sauer and Henderson, 1988; Sternberg and Hamilton, 1981) to retain compatibility with several mouse lines which rely on Cre/lox recombination (Kim et al., 2018). In order to stop nearby DNA elements from affecting expression of our constructs, we flanked both landing pads with full-length cHS4 insulator elements (Chung et al., 1993; Chung et al., 1997).

The first landing pad (“mKate2 landing pad”) (Fig. 1B) contains a splice acceptor and the *Neo* (G418/geneticin resistance) gene, the expression of which is driven by the *Rosa26* promoter and is thus dependent on correct targeting into intron 1. This cassette is followed by a cHS4 insulator, and a strong CAG promoter (Niwa et al., 1991) driving expression of the *mKate2-3xNLS* gene, which encodes a nuclear-localised red fluorescent protein with no evident phenotypic effect on mouse embryos (Malaguti et al., 2013; Shcherbo et al., 2009). The whole construct is flanked by attP50 SSR sites, which are recognised by the phiC31 integrase SSR (Huang et al., 2009), and followed by a second cHS4 insulator element.

The second landing pad (“EGFP landing pad”) (Fig. 1C) contains a splice acceptor and the *Hph* (hygromycin resistance) gene, the expression of which is driven by the *Rosa26* promoter and is thus dependent on correct targeting into intron 1. This cassette is flanked by Vlox2272 SSR sites, which are recognised by the VCre SSR (Suzuki and Nakayama, 2011), allowing its excision from the genome following targeting.

A VloxP site is present at the end of the landing pad, allowing for VCre-mediated exchange of the entire construct with a Vlox2272/VloxP-flanked donor vector. We flanked the rest of the landing pad with two cHS4 insulator sequences. The 5’ cHS4 insulator is followed by a roxP site, and the 3’ cHS4 insulator is preceded by a rox12 degenerate site, which are recognised by the Dre SSR (Chuang et al., 2016; Sauer and McDermott, 2004), allowing for Dre-mediated exchange of any cassette into this cHS4-insulated *Rosa26* locus. The roxP site is followed by a strong CAG promoter driving expression of *EGFP-3xNLS* (nuclear green fluorescent protein) followed by an internal ribosome entry site (IRES) and a *Ble* (bleomycin/zeocin resistance) gene. This *EGFP-3xNLS-IRES-Ble* cassette is flanked by FRT5 and FRT3 sites, which are recognised by the FLP SSR (Andrews et al., 1985; Schlake and Bode, 1994), allowing for exchange of this cassette with any FRT5/FRT3-flanked promoterless transgene, the expression of which would be driven by the upstream CAG promoter.

We targeted our landing pad constructs to the *Rosa26* locus, and obtained three clones with correct integration of both landing pads (Fig. S1A,B). Following VCre-mediated excision of the Vlox2272-flanked *Hph* resistance gene, we verified one clone had a normal diploid karyotype (Fig. S1C). We called this clonal cell line KELPE (knock-in exchangeable landing pad embryonic stem cells), after the Kelpie, a Scottish shape-shifting spirit who often appears in the shape of a horse (Burns, 1786; Gregor, 1883).

### Characterisation of KELPE PSCs

We set out to investigate whether the mKate2-3xNLS and EGFP-3xNLS transgenes were expressed by KELPE cells. We first tested whether both fluorescent proteins could be visualised by live imaging, by culturing KELPE cells on a Kelpie-shaped micropattern. Both mKate2-3xNLS and EGFP-3xNLS were expressed by all cells in culture, with a pattern consistent with nuclear localisation (Fig. 1D).

We next tested the ability of our transgenes to remain expressed following long-term pluripotent culture and long-term multilineage differentiation. We cultured KELPE cells for a minimum of 28 days in the absence of antibiotic selection in three conditions: ESC culture; LIF withdrawal, which drives multilineage differentiation (Morgani et al., 2013); and neural stem cell (NSC) differentiation (Pollard et al., 2006). We observed that both mKate2 and EGFP remained highly expressed in >99% cells across all three conditions, confirming our landing pads confer long-term protection from transgene silencing (Fig. 1E, Fig. S2).

### Design and assembly of donor vectors for RMCE

To generate donor vectors for RMCE into the KELPE landing pads, we designed a strategy based on EMMA cloning (Jones et al., 2019; Martella et al., 2017). This is a one-pot DNA assembly method based on Golden Gate cloning, optimised for complex mammalian expression vectors. It allows to assemble up to 25 functional modules (“parts”) in 25 pre-determined positions using 26 pre-tested fusion sites (A-Z) (Fig. S3). We generated a new library of parts by domesticating relevant functional units into Level 0 vectors (Fig. S3). Parts spanning different positions can be used to eliminate the need to use 25 Level 0 vectors for each assembly. Multiple functional units can be domesticated as a single part (Fig. 2A).

**Figure 2.**
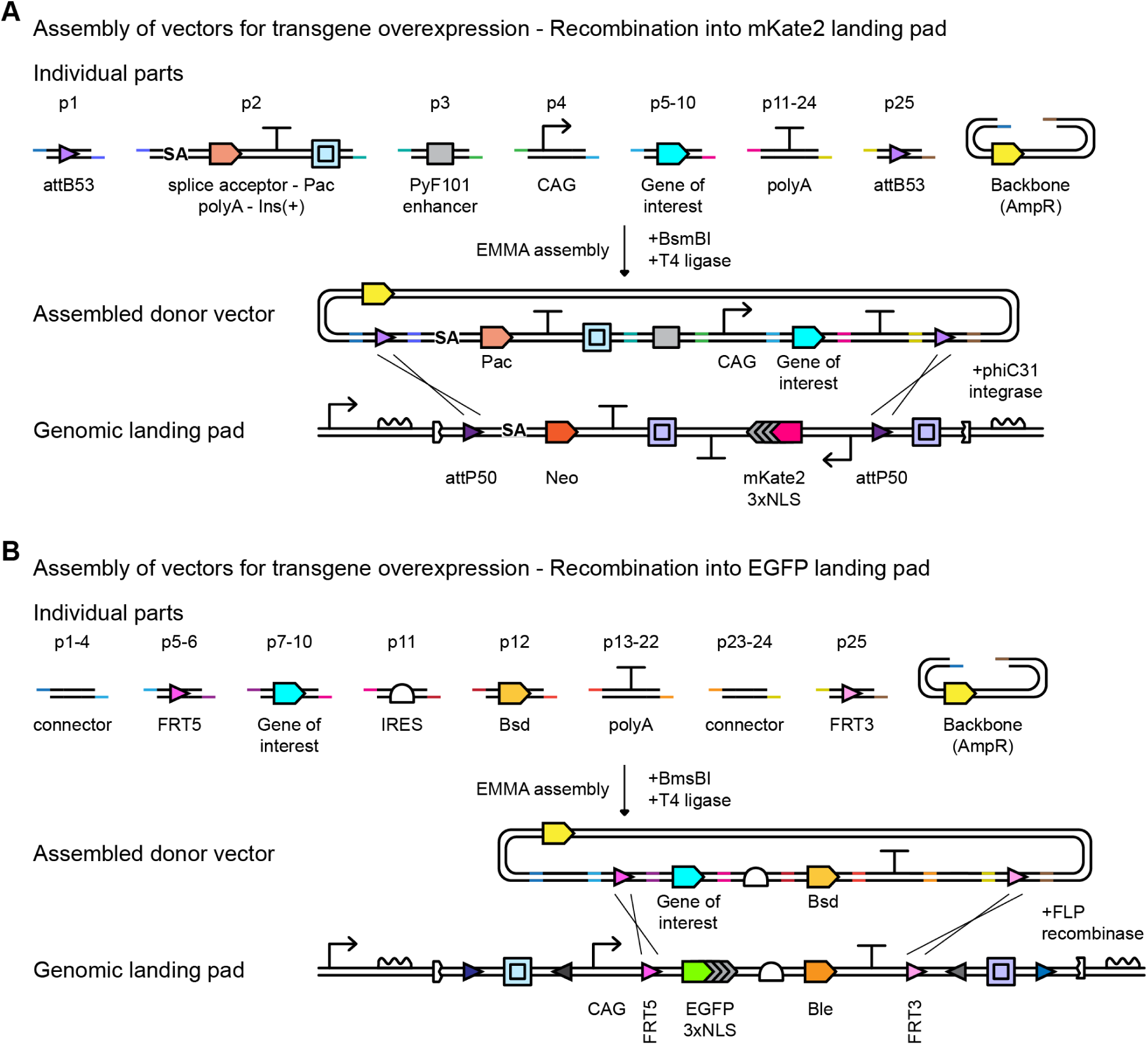
EMMA cloning strategy to assemble RMCE donor vectors for transgene overexpression. (A) Strategy to assemble donor vectors for transgene overexpression following RMCE into the mKate2 landing pad. The diagram illustrates Individual parts and positions within the EMMA assembly, the final assembled donor vector, and the RMCE event mediated by phiC31 integrase. Pac: puromycin N-acetyltransferase; AmpR: ampicillin resistance gene (beta-lactamase).sarala(B) Strategy to assemble donor vectors for transgene overexpression following RMCE into the EGFP landing pad. The diagram illustrates Individual parts and positions within the EMMA assembly, the final assembled donor vector, and the RMCE event mediated by FLP recombinase. Bsd: blasticidin S deaminase.

To build RMCE vectors for the mKate2 landing pad, we designed parts that when assembled would generate a donor vector containing attB53 sites (which can recombine with the attP50 sites in the landing pad) flanking a splice acceptor followed by the *Pac* (puromycin resistance) gene. This cassette is followed by a cHS4 insulator, a PyF101 enhancer and a CAG promoter driving expression of any transgene of interest, and a polyadenylation signal sequence. Correct phiC31 integrase-mediated RMCE would result in loss of G418/geneticin resistance, loss of mKate2-3xNLS fluorescence, and gain of puromycin resistance (Fig. 2A).

To build RMCE vectors for transgene overexpression from the EGFP landing pad, we designed parts that when assembled would generate a small promoterless donor vector containing FRT5 and FRT3 sites flanking any transgene of interest, an IRES, a *Bsd* (blasticidin resistance) gene and a polyadenylation signal sequence. Correct FLP-mediated recombination would result in loss of bleomycin/zeocin resistance, loss of EGFP-3xNLS fluorescence, and gain of blasticidin resistance (Fig. 2B).

### Screening of PUFFFIN neighbour-labelling transgenes

We next set out to illustrate how KELPE cells can be used to effectively screen a series of similar transgenes. We recently developed a synthetic neighbour-labelling system called PUFFFIN (Positive Ultrabright Fluorescent Fusion For Identifying Neighbours) (Lebek et al., 2024). This technology enables the identification of neighbours of “secretor” cells of interest. Secretor cells are engineered to express a nuclear fluorescent protein (to distinguish them from wild-type neighbours) and a secreted highly positively charged version of superfolder GFP (s36GFP). The high positive charge causes s36GFP to interact with the negatively charged plasma membranes of neighbouring cells, which enables uptake of s36GFP via endocytosis (Lebek et al., 2024; McNaughton et al., 2009; Thompson et al., 2012). s36GFP is very weakly fluorescent, and labelled neighbours cannot be detected by flow cytometry unless s36GFP is linked to a bright signal amplifier, such as mNeonGreen or HaloTag (Lebek et al., 2024). HaloTag is a modified haloalkane dehydrogenase that can covalently bind a range of “HaloTag ligands”, such as the fluorogenic far red Janelia Fluor 646 HaloTag Ligand (“JF646”) (Grimm et al., 2015; Los et al., 2008). We termed the s36GFP-HaloTag fusion “PUFFHalo” (Lebek et al., 2024) (Fig. 3A).

**Figure 3.**
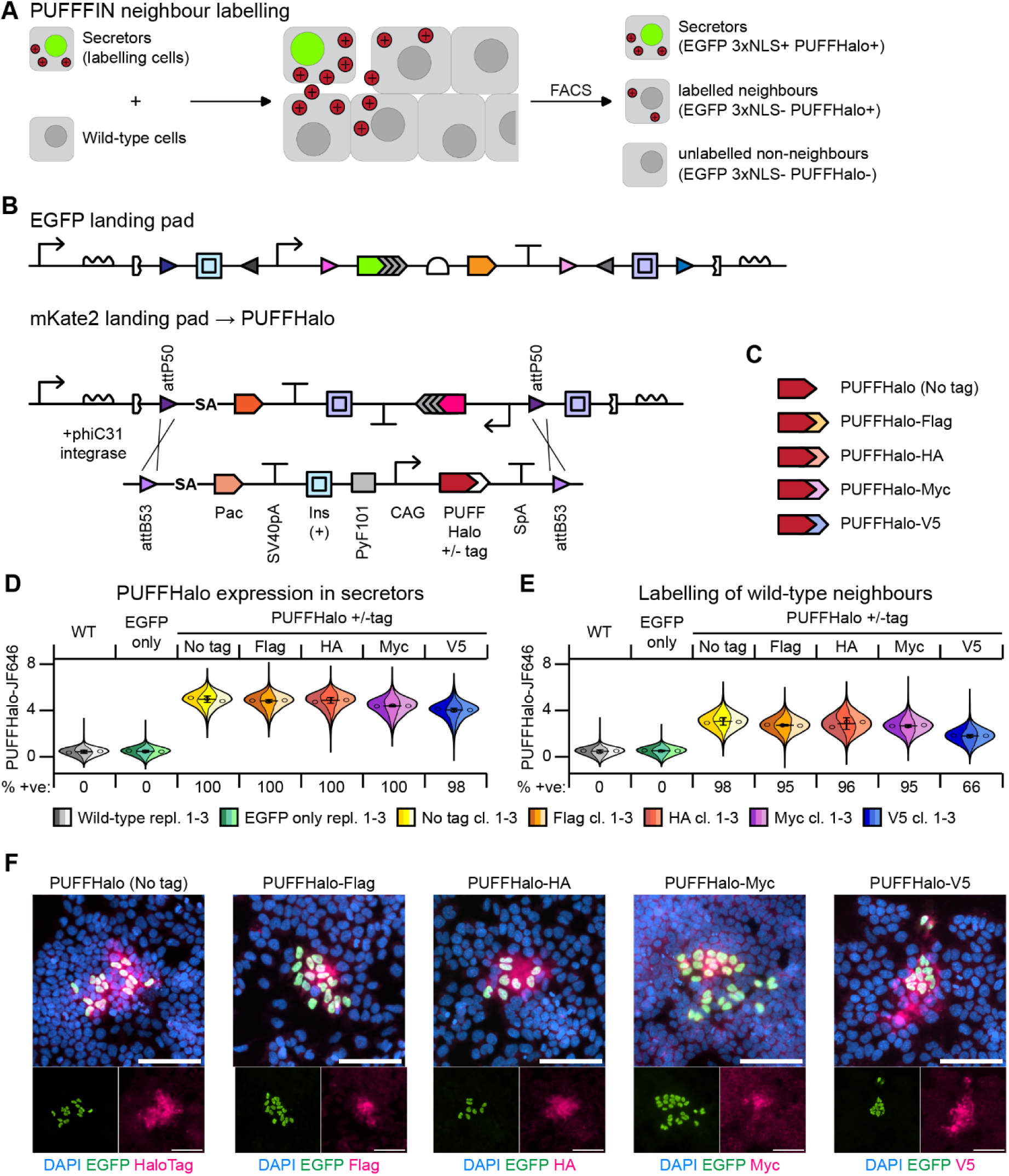
Screening of PUFFHalo constructs. (A) Diagram of PUFFFIN neighbour labelling. Secretor cells express nuclear EGFP-3xNLS and secrete a positively charged PUFFHalo label (red circle), which is uptaken by wild-type neighbouring cells. Flow cytometry allows to distinguish EGFP+ PUFFHalo+ secretors, EGFP-PUFFHalo+ neighbours of secretors, and EGFP-PUFFHalo-non-neighbours. (B) RMCE strategy to exchange a PUFFHalo overexpression cassette into the mKate2 landing pad. Pac: puromycin N-acetyltransferase; SV40pA: SV40 polyadenylation signal sequence. Ins: cHS4 insulator; PyF101: PyF101 enhancer; SpA: synthetic polyadenylation signal sequence. (C) Different PUFFHalo transgenes used for RMCE. (D) Flow cytometry analysis of PUFFHalo fluorescence in secretor cells stained with Janelia Fluor 646 HaloTag Ligand (“JF646”). (E) Flow cytometry analysis of PUFFHalo fluorescence in wild-type cells co-cultured with secretor cells (19:1 secretor:wild-type ratio) for 48 hours, stained with JF646. WT: wild-type; EGFP only: cell line containing EGFP landing pad but not PUFFHalo transgene. Violin Superplots display three individual replicates (WT, EGFP only) or three independent clones (all PUFFHalo lines) in different shading within the same distribution. Central line: median; error bars: 95% confidence interval. Nine-thousand cells were analysed for each replicate. (F) lmmunofluorescence of 1:99 secretor:wild-type 48-hour co-cultures, stained with antibodies indicated in figure. Scale bars: 100 µm.

There are few commercially available anti-HaloTag antibodies that have been cited in peer-reviewed literature, all raised in either mouse or rabbit, limiting potential co-staining applications. We therefore set out to test whether we could tag the C-terminus of PUFFHalo with commonly used protein tags recognised by multiple commercially available antibodies, and whether this would affect PUFFHalo expression and/or its ability to label neighbours.

We generated new clonal PUFFHalo secretor lines using KELPE cells. We retained the EGFP-3xNLS landing pad as a secretor cell marker, and replaced the mKate2 landing pad with a PUFFHalo overexpression cassette encoding either untagged PUFFHalo or PUFFHalo tagged with Flag, HA, Myc or V5 C-terminal tags (Fig. 3B,C).

We measured PUFFHalo and EGFP expression in secretor cells by flow cytometry, and observed that all lines retained high and similar levels of PUFFHalo and EGFP expression (Fig. 3D, Fig. S4A). We then tested neighbour-labelling efficiency by co-culturing a 19:1 ratio of secretors to wild-type cells, to ensure every wild-type cell was neighbouring at least one secretor cell, and measured the levels of PUFFHalo label in wild-type cells by flow cytometry. Whilst untagged PUFFHalo and Flag-, HA- and Myc-tagged PUFFHalo all labelled wild-type neighbours with similar high efficiency (>95%), the efficiency of labelling for PUFFHalo-V5 was lower (66%) (Fig. 3E, Fig. S4B-D), suggesting that a C-terminal V5 tag negatively affects the ability of PUFFHalo to label neighbours.

We next tested whether we could detect PUFFHalo by immunofluorescence in neighbours of secretor cells, by co-culturing a 1:99 ratio of secretor to wild-type cells, a ratio at which the majority of wild-type cells remains unlabelled (Fig. S4E). We could detect clear PUFFHalo signal in the vicinity of secretor cells using anti-HaloTag, anti-Flag, anti-HA and anti-V5 antibodies, but the signal was barely above background when using an anti-Myc antibody (Fig. 3F). The anti-Myc antibody stained a positive control very clearly (Fig. S4F), suggesting that the Myc epitope is poorly accessible when fused to the C-terminus of PUFFHalo.

Overall, we demonstrated that PUFFHalo, PUFFHalo-Flag and PUFFHalo-HA have extremely high neighbour-labelling efficiency (>95%), and that they can be detected by immunofluorescence with anti-HaloTag, anti-Flag and anti-HA antibodies.

This set of experiments demonstrated how KELPE cells can be used to perform rigorous and reliable comparisons of constructs expressed from the same genomic locus in sister cell lines.

### Generation of non-leaky SyNPL neighbour-labelling receiver cells

PUFFFIN allows users to identify cells in the vicinity of secretors, but does not allow to discriminate between direct neighbours and nearby cells (Lebek et al., 2024). We previously described SyNPL (Synthetic Notch Pluripotent cell lines), an adaptation of synthetic Notch (synNotch) technology (Morsut et al., 2016), that allows users to identify and manipulate direct neighbours of a cell of interest (Malaguti et al., 2022). In this system, a “sender” cell of interest is engineered to express a synthetic signal, membrane-tethered extracellular EGFP. “Receiver” cells are engineered to express a nuclear blue fluorescent protein (tagBFP-3xNLS) to facilitate their identification, an inducible fluorescent reporter cassette (*TRE-mCherry*) and an anti-EGFP synNotch receptor. This receptor comprises an extracellular anti-EGFP nanobody, the mouse Notch1 transmembrane core, and an intracellular tTA (tetracycline transactivator) synthetic transcription factor. The Notch1 core consists of a negative regulatory region (NRR), S2 and S3 protease cleavage sites, and a transmembrane domain. In the absence of receptor-ligand interaction, the NRR shields the S2 and S3 cleavage sites from endogenous proteases. Exertion of mechanical tension on the receptor due to receptor-ligand interaction displaces the NRR, leading to receptor cleavage and release of the tTA. The tTA can then translocate into the nucleus, bind to the TRE promoter and drive expression of the *mCherry* red fluorescent reporter (Malaguti et al., 2022) (Fig.4A).

**Figure 4.**
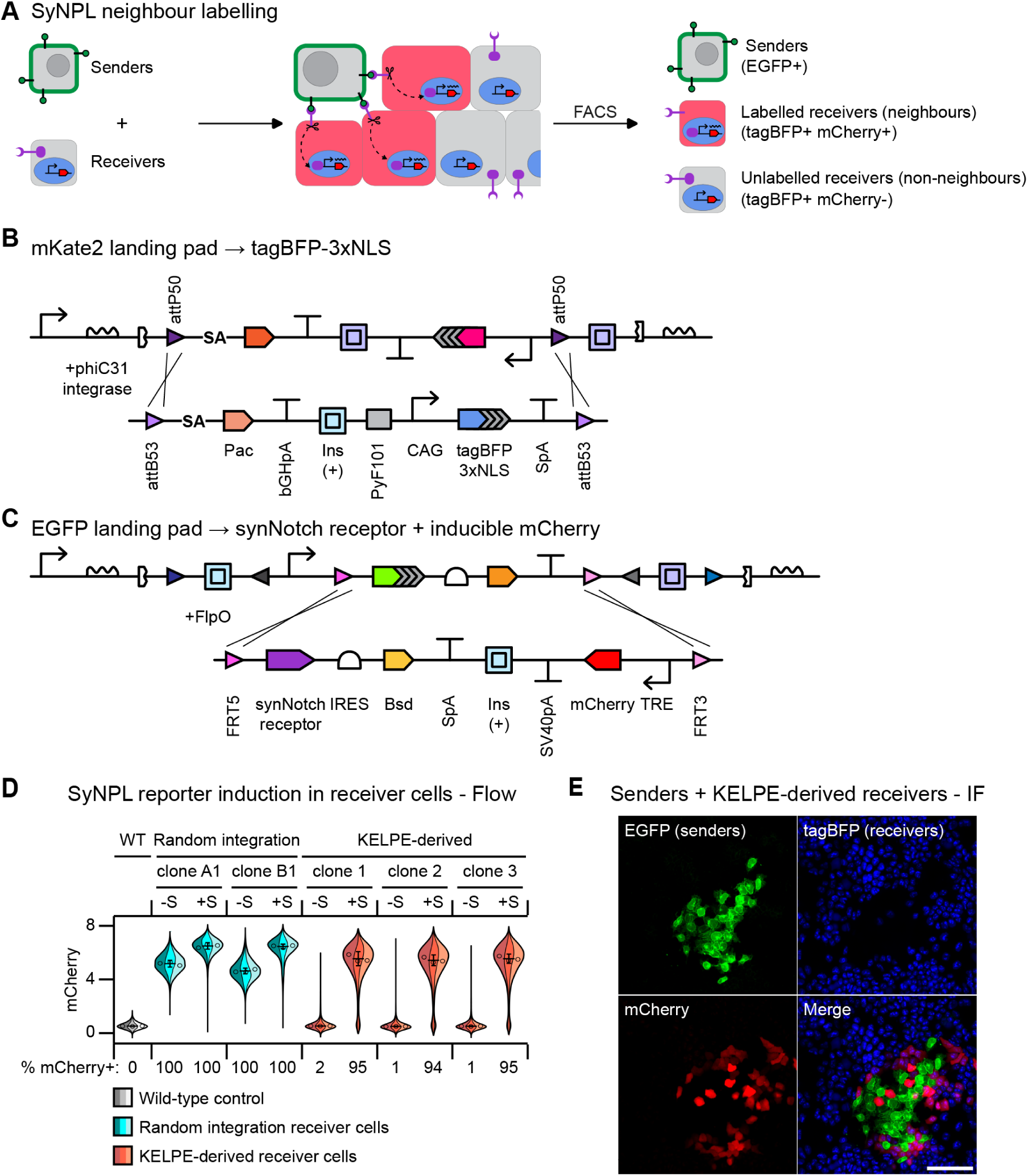
Generation of new SyNPL receiver cell lines. (A) Diagram of SyNPL neighbour labelling. Sender cells express plasma membrane-tethered extracellular EGFP. Receiver cells contain an inducible TRE-mCherry cassette, and express a nuclear tagBFP-3xNLS and a chimaeric synNotch receptor. The synNotch receptor comprises an anti-EGFP nanobody (LaG17), the mouse Notch1 core, and an intracellular tTA. Upon interaction between EGFP on sender cells and the synNotch receptor on receiver cells, the tTA is released from the receptor, translocates into the nucleus, binds TRE and induces mCherry expression. Flow cytometry allows to distinguish EGFP+ sender cells, mCherry+ tagBFP+ neighbours of sender cells, and mCherry- tagBFP+ non-neighbours. (B) RMCE strategy to exchange a tagBFP-3xNLS overexpression cassette into the mKate2 landing pad. (C) RMCE strategy to exchange a synNotch receptor expression cassette and a TRE-mCherry inducible cassette into the EGFP landing pad. (D) Flow cytometry analysis of mCherry fluorescence in wild-type cells (WT), previously published “random integration” SyNPL receiver cells, and KELPE-derived SyNPL receiver cells alone (“-S”) or in co-culture with sender cells for 48 hours (“+S”) (19:1 sender:receiver ratio). Violin Superplots display three independent replicates in different shading within the same distribution. Central line: median; error bars: 95% confidence interval. Ten-thousand cells were analysed for each replicate. (E) lmmunofluorescence of 1:99 sender:receiver 48-hour co-cultures, stained with antibodies indicated in figure. Scale bar: 100 µm.

We previously generated SyNPL receiver cells by random genomic integration of the anti-EGFP synNotch receptor and *tagBFP-3xNLS* constructs, and RMCE of the *TRE-mCherry* cassette into a non-insulated *Rosa26* landing pad. These receiver cells displayed a clear increase in mCherry fluorescence following co-culture with sender cells, but also had baseline mCherry leakiness when cultured alone (Malaguti et al., 2022; Santorelli et al., 2024). Whilst this does not affect the capacity of distinguishing mCherry-high neighbours from mCherry-low non-neighbours, it might prove problematic for users wishing to express functional transgenes in neighbours of sender cells of interest.

Interestingly, the baseline mCherry leakiness in these cells could not be abolished by inhibition of synNotch receptor cleavage nor with inhibition of tTA DNA binding (Malaguti et al., 2022), suggesting that enhancers near the *Rosa26-TRE-mCherry* cassette may be responsible for baseline *mCherry* expression.

We set out to test whether we could leverage the cHS4 insulator elements in the KELPE lines to generate new receiver lines with no baseline mCherry leakiness. We replaced the mKate2 landing pad with a *PyCAG-tagBFP-3xNLS* cassette (Fig. 4B), and the sequence downstream of the CAG promoter in the EGFP landing pad with the anti-EGFP synNotch receptor transgene, a cHS4 insulator sequence, and a *TRE-mCherry* cassette (Fig. 4C).

We performed flow cytometry on three KELPE-derived receiver cell clones and confirmed they expressed high and uniform levels of tagBFP, outperforming our previous random integration receiver cells (Fig. S5A-C). We then compared mCherry levels in our previous random integration receiver cells and our new KELPE-derived receiver cells. We did so by culturing receiver cells alone or with an excess of sender cells (19:1 sender:receiver ratio), to ensure each receiver cell made contact with sender cells (Fig. S5D). We observed that unlike random integration receiver cells, KELPE-derived receiver cells had extremely low (1-2%) mCherry leakiness in the absence of sender cells, whilst retaining high inducibility (>94%) in the presence of sender cells (Fig. 4D, Fig. S5E). This suggests that insulation of the *Rosa26* locus allows for much tighter regulation of the TRE inducible promoter. We also observed that all syngeneic KELPE-derived clones behaved virtually identically across all assays, as expected (Fig. 4D, Fig. S5C).

Finally, we tested whether sender cells could specifically label KELPE-derived receiver neighbours, by co-culturing a 1:99 ratio of sender:receiver cells. We observed mCherry label expression in neighbours of sender cells (Fig. 4E), suggesting that KELPE-derived receiver cells can be used to identify interactions with sender cells of interest.

Overall, we demonstrated how the insulator elements in KELPE cells can shield transgenic cassettes from the influence of endogenous DNA regulatory elements, and generated new and improved SyNPL receiver clonal cell lines.

### Screening of plasma membrane-tethered EGFP transgenes

The SyNPL sender cells we previously reported were generated via random integration of a construct encoding EGFP fused to the human PDGFRB transmembrane domain (“EGFP-TMD”) (Malaguti et al., 2022). This fusion protein had previously been shown to efficiently activate anti-EGFP synNotch receptor cleavage in mammalian cell lines (Morsut et al., 2016). EGFP can also be delivered to the outer leaflet of the plasma membrane by fusing it to a GPI-lation peptide (“EGFP-GPI”) (Hiscox et al., 2002). Several EGFP-GPI-expressing mouse lines have been independently generated, with no overt phenotype (Kondoh et al., 1999; Rhee et al., 2006; Wiese et al., 2013), suggesting EGFP-GPI may serve as a useful alternative to EGFP-TMD if it were able to induce reporter activation in SyNPL receiver cells.

We set out to perform a rigorous comparison of EGFP-TMD and EGFP-GPI by integrating each construct into the KELPE EGFP landing pad in place of EGFP-3xNLS (Fig. 5A). We compared expression of EGFP-3xNLS, EGFP-TMD, and EGFP-GPI in pluripotency culture conditions and after 48 hours of differentiation in N2B27, a time frame that allows mESCs to exit naïve pluripotency (Betschinger et al., 2013; Kalkan et al., 2017; Malaguti et al., 2013). The pattern of EGFP expression was consistent with nuclear (EGFP-3xNLS) or plasma membrane (EGFP-TMD, EGFP-GPI) localisation in both pluripotency and N2B27 differentiation (Fig. 5B). We measured levels of EGFP by flow cytometry, and observed highest fluorescence in EGFP-3xNLS-expressing cells, followed by EGFP-GPI, with EGFP-TMD displaying lowest fluorescence (Fig. 5C), consistent with immunofluorescence data (Fig. 5B). Fluorescence levels were comparable between pluripotency and N2B27 differentiation (Fig. 5C), which is consistent with the lack of EGFP silencing observed in long-term differentiation of parental KELPE cells (Fig. 1E).

**Figure 5.**
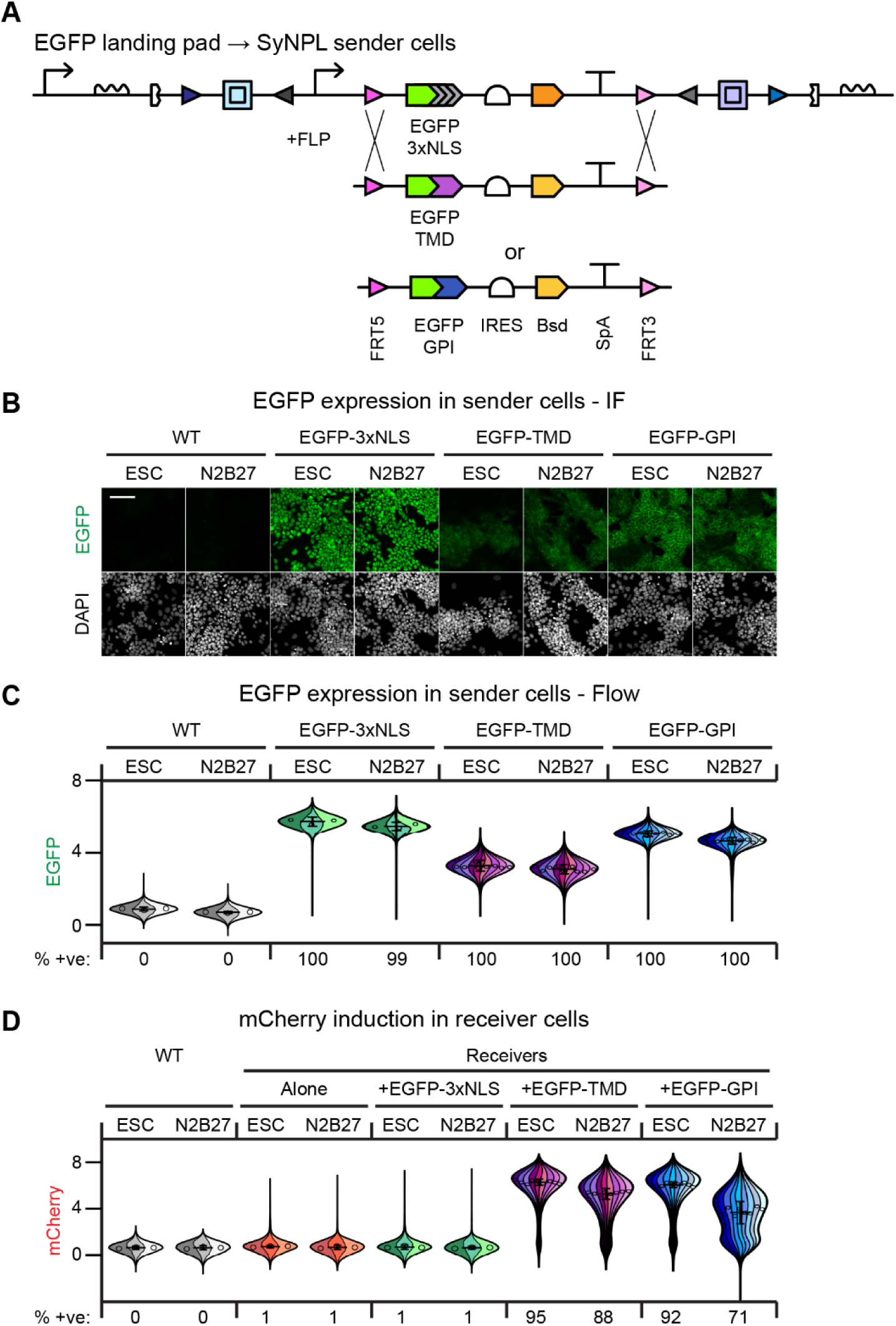
Screening of plasma membrane-tethered EGFP transgenes. (A) RMCE strategy to exchange EGFP-TMD or EGFP-GPI into the EGFP-3xNLS landing pad.sarala(B-C) lmmunofluorescence (B) or flow cytometry analysis (C) of EGFP expression in wild-type (“WT”) cells or cells expressing EGFP-3xNLS, EGFP-TMD or EGFP-GPI from the KELPE EGFP landing pad, in pluripotency culture conditions (“ESC”) or after 48 hours in N2B27 differentiation medium (“N2B27’’). Scale bar: 100 μm.sarala(D) Flow cytometry analysis of mCherry fluorescence in wild-type cells and KELPE-derived SyNPL receiver cells cultured alone or following 48 hours of co-culture (19:1 sender:receiver ratio) with KELPE-derived SyNPL sender cells expressing EGFP-3xNLS (negative control), EGFP-TMD or EGFP-GPI, in ESC and N2B27 conditions. Violin Superplots display three independent replicates for one (WT, Receivers Alone, EGFP-3xNLS) or three (EGFP-TMD, EGFP-GPI) independent clones in different shading within the same distribution. Central line: median; error bars: 95% confidence interval. Ten-thousand cells were analysed for each replicate in (C), six-thousand five-hundred cells were analysed for each replicate in (D).

We next tested the ability of each cell line to induce *mCherry* expression in KELPE-derived SyNPL receiver cells, performing flow cytometry following a 19:1 sender:receiver co-culture, and measuring mCherry levels in receiver cells (Fig. 5D). As expected, receiver cells cultured alone were mostly negative for mCherry fluorescence (1% positive cells). Co-culture of receiver cells with “mock sender” cells expressing EGFP-3xNLS also resulted in no mCherry upregulation. In pluripotency culture conditions, EGFP-TMD and EGFP-GPI performed comparably, with the majority (>92%) of receiver cells inducing mCherry (Fig. 5D). This suggests either that anti-EGFP synNotch receptors on receiver cells can be fully saturated by the lower levels of EGFP observed in EGFP-TMD cells, or that activation of a subset of anti-EGFP synNotch receptors may be sufficient for maximum *mCherry* transcriptional output. Strikingly, EGFP-GPI-expressing cells were comprehensively outperformed by EGFP-TMD-expressing cells in N2B27 differentiation, with a drastic reduction in mCherry fluorescence observed in co-cultures with EGFP-GPI sender cells (Fig. 5D). This result was surprising, as the levels of EGFP-GPI in N2B27 are comparable to those observed in pluripotency culture conditions (Fig. 5C).

Overall, we demonstrated that EGFP-GPI can act as a ligand for anti-EGFP synNotch receptors, that it is expressed at higher levels than EGFP-TMD, and that it is able to induce mCherry expression in receiver cells in pluripotency culture conditions, but fails to drive mCherry expression in a subset of receiver cells following 48 hours of N2B27 differentiation. By comparing transgenes expressed from the same genomic locus in congenic cell lines, we were able to draw this unexpected conclusion following a single set of screening experiments, illustrating the usefulness of KELPE cells for genetic circuit prototyping.

### Synthetic programming of contact-induced cell death

Having leveraged KELPE cells to generate new optimised SyNPL receiver and sender cells, we set out to showcase the opportunities conferred by our new low-leakiness SyNPL receiver transgene structure. The SyNPL system allows users to engineer receiver cells so that they express any transgene following interaction with sender cells. This can be a reporter, to monitor cell interactions (Fig. 4), or a functional transgene, to manipulate the behaviour of receiver cells following interaction with sender cells (Malaguti et al., 2022).

We attempted to generate receiver cells capable of inducing the toxic peptide DTA (diphtheria toxin fragment A). DTA contains the catalytic subunit of diphtheria toxin (Pappenheimer, 1977), and it is commonly used in mouse PSCs as a negative selection marker (Chu et al., 2004; Malaguti et al., 2019; Tessarollo et al., 2009). A single DTA molecule is capable of killing mammalian cells (Yamaizumi et al., 1978), implying that in order for inducible DTA receiver cells to survive, they must not express basal levels of the *DTA* transgene in the absence of sender cells. We adopted the same RMCE strategy we used to generate inducible mCherry receivers, replacing the *mCherry* transgene with a *DTA* transgene in our donor vector (Fig. 6A,B). We obtained EGFP-negative blasticidin-resistant clonal cell lines after RMCE, suggesting that the lines had negligible leakiness. We next set out to test whether we could induce DTA-mediated killing of receiver cells by co-culturing inducible DTA receiver cells with an excess of sender cells (19:1 sender:receiver ratio). As a control, we performed the same co-cultures in the presence of doxycycline, a cell-permeable antibiotic that can bind the tTA element of the synNotch receptor, prevent it from binding DNA and thus stop it from driving transcription of *DTA* from the TRE inducible promoter (Courte et al., 2024; Malaguti et al., 2022) (Fig. 6C). As an additional control, we performed the same experiment with receiver cell lines that did not carry an inducible DTA cassette. Following co-culture, we observed fewer receiver cells in DTA-inducible receiver cells co-cultured with sender cells in the absence of doxycycline than in any of the control conditions (Fig. 6D, Fig. S6). To quantify this observation, we performed flow cytometry to calculate the proportion of receiver cells in co-cultures without doxycycline over cultures with doxycycline (Fig. 6E). Should DTA induction have no effect, we would expect a ratio of 1, and should DTA kill all receiver cells, we would expect a ratio of 0. We observed ratios of approximately 1 for three control (no DTA) receiver cell clones, and ratios of 0.05-0.07 for three inducible DTA receiver cell clones (Fig. 6E). This confirms that sender cells can induce *DTA* expression in receiver cells following their interaction, and that DTA is capable of killing receiver cells.

**Figure 6.**
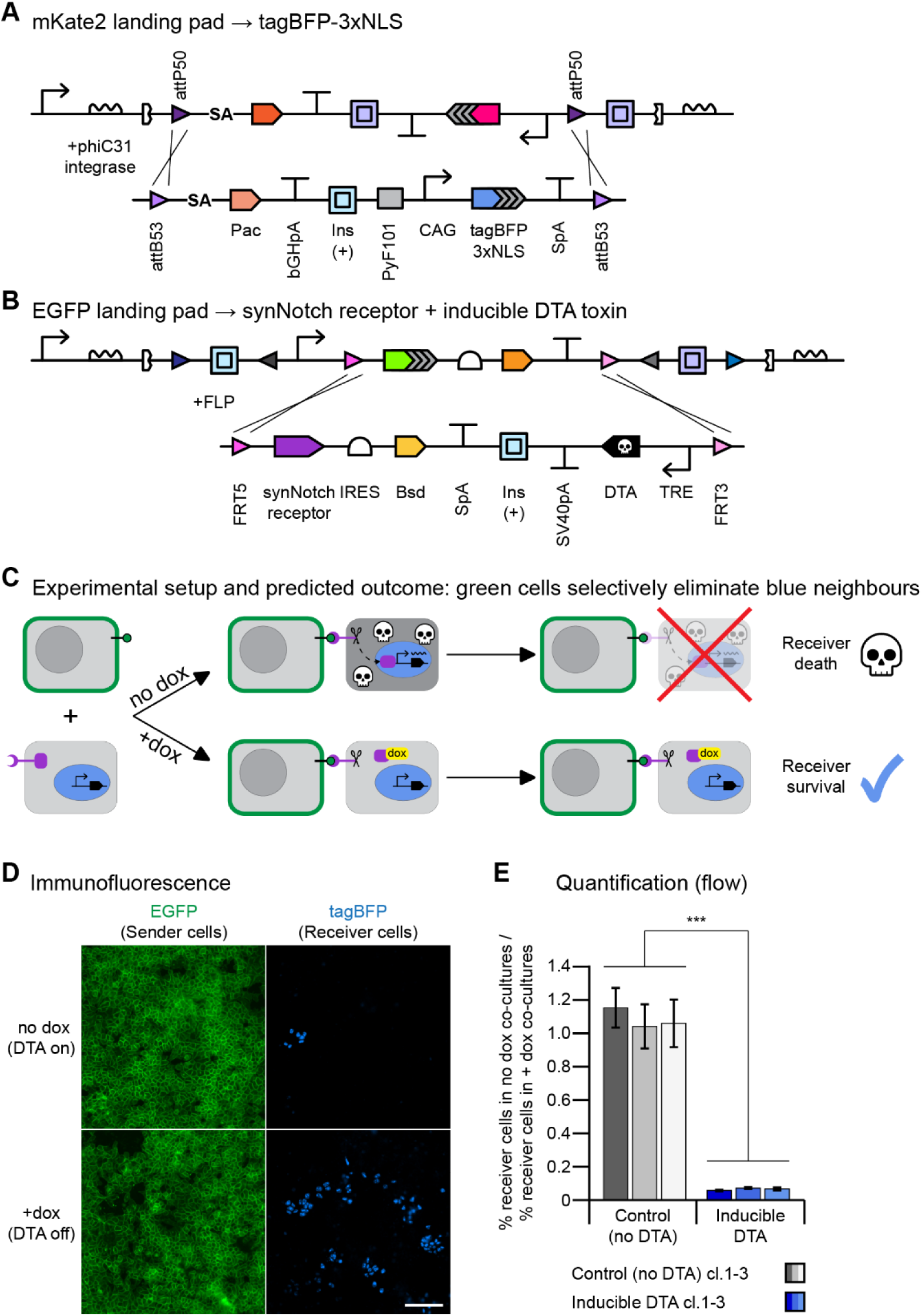
Generation of inducible DTA toxin SyNPL receiver cells. (A) RMCE strategy to exchange a tagBFP-3xNLS overexpression cassette into the mKate2 landing pad. (B) RMCE strategy to exchange a synNotch receptor expression cassette and a TRE-DTA inducible cassette into the EGFP landing pad. (C) Diagram illustrating the experimental setup and predicted outcome. EGFP+ sender cells were co-cultured with tagBFP+ receiver cells (19:1 sender:receiver co-culture) in the presence or the absence of doxycycline, which inhibits tTA from binding TRE and driving OTA expression. In the absence of doxycycline, OTA expression should lead to death of receiver cells. In the presence of doxycyline, receiver cells should survive. (D) lmmunofluorescence for EGFP and tagBFP in 19:1 sender:receiver cell co-cultures in the absence (“no dox”) or presence (“+dox”) of doxycycline. Scale bar: 100 µm. (E) Flow cytometry quantification of the ratio of receiver cells found in “no dox” over “+dox” co-cultures, in control cells lacking an inducible OTA transgene and in inducible OTA SyNPL receiver cells. A ratio of 1 indicates no difference between “no dox” and “+dox” conditions. Bars: mean of three biological replicates for each of three independent clones for each cell line. Error bars: standard deviation. Statistical test: nested t-test; *** p<0.001.

Overall, this set of experiments demonstrates that KELPE-derived SyNPL receiver cells have negligible leakiness, and suggests that they are suitable for expression of any functional transgene, including toxins. These data also illustrate how this KELPE-derived SyNPL system can be used to synthetically program cell death, and how the process can be externally controlled via administration of doxycyline.

## DISCUSSION

The design, prototyping and establishment of genetic circuitry in mouse pluripotent stem cells is a challenging process. Here, we describe the first mESC line carrying two *Rosa26*-targeted insulated genomic landing pads, illustrate how it allows for the convenient generation of silencing-resistant multi-transgene-expressing cell lines, and demonstrate how it can be used to rigorously screen different synthetic circuit components.

### Choice of site-specific recombinase sites

Many SSR sites have been described and tested in mammalian cells. Due to our previous experience using phiC31/att and FLP/FRT RMCE systems (Malaguti et al., 2019; Malaguti et al., 2022), we included these sites in our landing pads. When generating the modular EGFP landing pad, we selected other SSR sites that had previously been shown to work in mESCs: Dre/rox (Anastassiadis et al., 2009) and VCre/Vlox (Suzuki and Nakayama, 2011). We chose to avoid using the Cre/lox SSR system to maintain our landing pads fully orthogonal with any mouse Cre-driver lines, or any mESCs containing Cre-ERT2 tamoxifen-inducible systems. Alternative SSR systems, such as SCre/Slox, Vika/vox or Bxb1/att, which have been shown to also be functional in PSCs or mice (Karimova et al., 2013; Karimova et al., 2018; Russell et al., 2006; Suzuki and Nakayama, 2011), could also be recombined into the KELPE landing pads to add further modularity to the system.

### Choice of promoters and polyA sequences

For transgene overexpression, we relied on the strong ubiquitously expressed CAG promoter (Niwa et al., 1991), which has been shown to drive higher levels of transgene overexpression than other promoters when integrated into the *Rosa26* locus (Chen et al., 2011a). For synNotch receptor-mediated transgene induction, we made use of a “TRE” promoter consisting of 7 tet-responsive elements followed by a minimal CMV promoter. We previously demonstrated how this promoter integrated into the *Rosa26* locus outperformed an analogous *Rosa26-*targeted “tetO” promoter with 10 tet-responsive elements followed by a minimal CMV promoter element (Malaguti et al., 2022). Finally, we used a unique set of polyadenylation signal sequences to mitigate the risk of mitotic recombination (Stern, 1936).

### PUFFFIN cell lines

We previously established random integration mESC PUFFHalo secretor cells, which exhibited a 1-log difference in levels of nuclear fluorescent protein and PUFFHalo label compared to background levels observed in wild-type cells by flow cytometry (Lebek et al., 2024). The KELPE-derived PUFFHalo secretor lines we describe here display a 2-log difference in levels of both EGFP-3xNLS and of PUFFHalo label compared to wild-type cells (Fig. S4B,E), a considerable improvement over our random integration cell lines, in which both PUFFHalo and the nuclear fluorescent protein were expressed monocistronically. This suggests that segregation of the two transgenes into separate expression cassettes can lead to higher expression of both proteins.

It is unclear why the C-terminal fusion of a V5 tag would affect PUFFHalo’s ability to label neighbouring cells. At pH 7.4, Flag, HA and Myc tags have a slight overall negative charge (−3, −2, and −3 respectively) whereas the V5 tag has an overall neutral charge (Fritschle et al., 2025). The V5 tag should therefore have the least influence on PUFFHalo’s ability to interact with negatively charged plasma membranes. It is thus likely that the V5 tag is affecting structural properties of PUFFHalo, which could in turn lead to reduced labelling of neighbouring cells.

### Comparison of membrane-tethered EGFP constructs

We demonstrated that EGFP-GPI can act as a ligand for anti-EGFP synNotch receptors. We observed that EGFP-GPI cells had higher EGFP fluorescence than those expressing EGFP-TMD in both pluripotent cultures and in N2B27 differentiation (Fig. 5C). However, EGFP-GPI-expressing sender cells performed worse than EGFP-TMD-expressing sender cells at inducing expression of an mCherry reporter in SyNPL receiver cells in N2B27 culture (Fig. 5D). Cleavage of anti-EGFP synNotch receptors requires mechanical tension exerted trough interaction of the synNotch receptor with plasma membrane-tethered EGFP (Garibyan et al., 2024; Gordon et al., 2015; Hamann et al., 2025; Morsut et al., 2016; Roybal et al., 2016; Sloas et al., 2023; Toda et al., 2020; Zhang et al., 2022). During exit from naïve pluripotency, plasma membrane tension decreases in mESCs (Bergert et al., 2021; De Belly et al., 2021). EGFP-GPI is anchored to the outer leaflet of the plasma membrane (Hiscox et al., 2002), whereas the PDGFRB transmembrane domain in EGFP-TMD spans the entire plasma membrane (Piñero-Lambea et al., 2015). It is therefore possible that during early differentiation, EGFP-GPI is no longer capable of exerting enough tensile strength on the anti-EGFP synNotch receptor to drive uniform mCherry reporter induction in all receiver cells.

### Non-leaky SyNPL receiver cell lines

Ligand-independent activation of synNotch receptors is a widely reported issue (Asaadi and Rahbarizadeh, 2025; He et al., 2017; Morsut et al., 2016; Semeniuk et al., 2024; Sloas et al., 2023; Wang et al., 2023; Williams et al., 2020; Yang et al., 2020). This is perhaps unsurprising, as endogenous Notch signalling is also known to exhibit ligand-independent activation (Palmer and Deng, 2015). We previously established SyNPL receiver cells via random integration of constructs encoding tagBFP-3xNLS and EGFP-Notch1 core-tTA synNotch receptor, and RMCE of *TRE-mCherry* into a non-insulated *Rosa26* landing pad (Malaguti et al., 2022). These cells exhibited leaky mCherry reporter expression in the absence of sender cells, but, interestingly, this leakiness could not be abolished by inhibiting tTA-induced mCherry transcription. This suggests that cellular mechanisms other than ligand-independent receptor activation must be at play. Here, we demonstrate that by insulating the *TRE-mCherry* cassette we can generate new SyNPL receiver cells with extremely low baseline mCherry expression (<2%) and high inducibility (>94%) (Fig. 4D). This suggests that in the context of non-insulated SyNPL receiver mESCs, the effect of nearby enhancer elements on the TRE promoter far outweighs the effect of ligand-independent synNotch receptor activation.

Furthermore, these observations suggest that *Rosa26*-targeted insulated “TRE-gene of interest” cassettes have negligible leakiness. It is therefore possible that these cassettes could be paired with constitutive expression of rtTA or tTA to generate doxycycline-inducible or doxycycline-repressible systems which may allow expression of toxic proteins without having to rely on complex genetic circuitry to abolish baseline expression (De Carluccio et al., 2024; Hosoda et al., 2011; Kiwimagi et al., 2021).

We expect that it will be possible to repurpose this synNotch receiver transgene syntax to tightly regulate the induction of any transgene of interest in mESCs, opening up huge opportunities to synthetically program cell behaviour in response to interaction with a particular cell of interest. We exemplified this by generating receiver cell lines in which we were able to synthetically program non-cell-autonomous contact-induced cell death.

It remains to be seen whether this low leakiness and high inducibility in mESCs is specific to EGFP ligand/anti-EGFP synNotch receptor pairings. An ALFA-tag/NbALFA synNotch system was recently established in a polyclonal mESC line, displaying approximately 40% inducibility (Soliman et al., 2025). RMCE-mediated integration of these constructs into KELPE cells would allow to test whether this synNotch receiver transgene syntax may be suitable for other ligand/synNotch receptor pairings.

### Concluding remarks

Transgenesis was first performed on mammalian cells half a century ago (Davidson et al., 1973; Munyon et al., 1971; Wigler et al., 1977). These pioneering studies identified key challenges: transgene silencing, variation in transgene expression, and efficiency of transgene delivery. Fifty years later, these same issues affect the development of synthetic gene circuits in mammalian cells (Alhaji et al., 2019; Cabrera et al., 2022).

Here, we demonstrate how KELPE cells allow users to overcome these issues, and provide a foundational platform for synthetic circuit development in mouse pluripotent stem cells.

## MATERIALS AND METHODS

### Reagents

All reagents are listed in Table S2.

### Mouse embryonic stem cell culture

Mouse embryonic stem cells were maintained on gelatinised culture vessels at 37°C and 5% CO2 in Glasgow Minimum Essential Medium (GMEM) supplemented with 10% foetal calf serum (FCS), 100 U/ml LIF (produced in-house), 0.5 µM PD0325901, 100 nM 2-mercaptoethanol, 1x non-essential amino acids, 2 mM L-glutamine and 1 mM sodium pyruvate. This medium was referred to as “ESC culture medium”. When appropriate, antibiotic selection were added as follows: 2 µg/ml puromycin, 200 µg/ml hygromycin B, 10 µg/ml blasticidin, 200 µg/ml G418. For passaging, cells were washed in Dulbecco’s phosphate buffered saline solution without calcium and magnesium (PBS), detached from their culture vessel using Trypsin-EDTA 0.05%, quenched in ESC culture medium, pelleted by spinning at 300g for 3 min, and resuspended in ESC culture medium, prior to replating at the desired density. Cells were passaged every 1-2 days. For live imaging, GMEM was replaced with Phenol Red-free Dulbecco’s Modified Eagle Medium (DMEM), with all other components of the culture medium used at identical concentrations. Cells were routinely tested to ensure absence of mycoplasma contamination.

### Differentiation assays

N2B27 medium was prepared as previously described (Pollard et al., 2006). Its composition is a 1:1 mixture of DMEM/F12 and Neurobasal medium, supplemented with 0.5X N-2 Supplement, 0.5X B-27 Supplement, 2mM L-Glutamine and 100nM 2-mercaptoethanol. For differentiation experiments, cells were trypsinised, washed twice in N2B27 and replated onto cell culture vessels coated with 7.5 µg/ml fibronectin.

“LIF withdrawal medium” was prepared using the same recipe as “ESC culture medium”, without adding LIF and PD0325901. To induce differentiation, cells were trypsinised, washed twice in LIF withdrawal medium, and replated onto gelatinised culture vessels. Differentiating cells were passaged in LIF withdrawal medium when required depending on cell density.

The differentiation of ESCs to NSCs was based on previously established protocols (Pollard, 2013; Pollard et al., 2006). Cells were passaged using Trypsin-EDTA 0.05% and washed twice with DMEM/F12. Cells were then plated at 1.25×10^5^ per well of a gelatine-coated 6-well plate in NSC media consisting of: DMEM/HAMS-F12, 8mM glucose, 1x non-essential amino acids, 0.012% Bovine Albumin Serum, 100µM 2-mercaptoethanol, 0.5X B-27 Supplement and 0.5X N-2 Supplement. Cells were allowed to differentiate for 7 days with daily media changes from day 2 onwards. On Day 7, cells were re-plated onto Laminin I-coated wells and from here on out cultured in NSC media with 10 ng/ml mEGF, 10 ng/ml hFGF2 and 2 µg/ml Cultrex Laminin I. They were then passaged every few days using Accutase onto Laminin I-coated culture vessels.

### EMMA assemblies

Functional DNA parts of interest were domesticated for EMMA assembly by amplifying them with primers with the following structure: N_6_-BsmBI binding site-N-fusion site-N_20_, using the fusion sites listed in the original EMMA publication (Martella et al., 2017). Alternatively, small DNA functional parts were custom synthesised with the same structure (N_6_-BsmBI binding site-N-5’ fusion site-DNA part-3’ fusion site-N-BsmBI binding site- N_6_). Domesticated parts were subcloned into pCR-BluntII-Topo vectors and sequence verified. EMMA assemblies were performed using the NEBridge® Golden Gate Assembly Kit (BsmBI-v2), using 75 ng of each DNA part and of the EMMA receiver vector (Addgene plasmid #100637, gift from Steven Pollard) (Martella et al., 2017), 2 µl T4 DNA ligase buffer, 2 µl BsmBI-v2+T4 DNA ligase enzyme mix, in a total volume of 20 µl. The reactions were subject to 30 cycles of 42°C for 5 minutes, 16°C for 5 minutes, followed by a 5 minute heat inactivation step at 60°C. 10 µl of the reaction was transformed into subcloning efficiency NEB® 5-alpha Competent *E. coli* or NEB® Stable Competent *E. coli* (high-efficiency) and plated onto ampicillin resistance Luria-Bertani broth agar plates. Colonies were picked, miniprepped, test digested, and clones with the expected banding pattern were sequenced verified.

### Generation of landing pad constructs

The mKate2 landing pad targeting construct was generated by modifying the pmROSA26-attP50-Neo-mKate2-3xNLS-attP50 plasmid (Addgene plasmid #183609) we previously described (Malaguti et al., 2022). We digested this plasmid with FspAI+AscI to remove an unnecessary bGHpA sequence located 3’ of the second attP50 site, and upstream of the *Rosa26* 3’ homology arm. We ligated this to an AscI-cHS4 insulator-EcoRV fragment generated by overlap extension PCR of the cHS4 insulator sequence from plasmid “GBSGFP insulator” (Addgene plasmid #121975, gift from James Briscoe) (Balaskas et al., 2012) followed by AscI+EcoRV digestion of the PCR amplicon. We replaced a PacI-NsiI fragment containing a splice acceptor-loxP-attB1-attP50-Neo-SV40pA-bGHpA with a custom synthesised PacI-NsiI fragment containing attP50-splice acceptor-Neo-SV40pA-cHS4 insulator-bGHpA to remove unwanted loxP and attB1 sites and introduce a second insulator sequence. This generated our final mKate2 landing pad plasmid pmROSA26-ANCKAC (attP50-Neo-cHS4-mKate2-attP50-cHS4).

The EGFP landing pad construct was generated *de novo* by EMMA assembly. A list of parts used is shown in Table S3. The landing pad construct was subcloned into the pmROSA26-ANCKAC targeting vector in place of the mKate2 landing pad, by digesting both vector and landing pad insert with PacI+AscI and ligating the two fragments together. This generated the EGFP landing pad plasmid pmROSA26-HELP (HygroR EGFP Landing Pad).

Due to their size and to the presence of repeated DNA elements (cHS4 insulators), pmROSA26-ANCKAC and pmROSA26-HELP were expanded in NEB® Stable Competent *E. coli* grown at 30°C.

### Generation of RMCE constructs

Constructs for RMCE into the mKate2 and EGFP landing pads were generated by EMMA assembly. A list of parts used for the various constructs is shown in Tables S4-S10. pCAG:GPI-GFP was a gift from Anna-Katerina Hadjantonakis (Addgene plasmid # 32601) (Rhee et al., 2006), and was used as a PCR template for domestication of the EGFP-GPI sequence.

### Generation of recombinase expression plasmids

Constructs to transiently express VCre and FLPo were generated by EMMA assembly. A list of parts used for the two constructs is shown in Tables S11-S12.

pCAG-VCre (Addgene plasmid # 89575) and pCAG-FlpO (Addgene plasmid # 89574) were gifts from Wilson Wong (Weinberg et al., 2017), and were used as PCR templates for domestication of the VCre and FLPo sequences.

The CAG-phiC31 integrase plasmid (Monetti et al., 2011) was a gift from Andras Nagy. An equivalent plasmid is available on Addgene (plasmid # 62658).

### Transfections

mESCs were nucleofected using the Lonza P3 Primary Cell 4D-Nucleofector X kit, program CG-104, as per manufacturer’s instructions. 5×10^5^ mESCs were nucleofected with 2-5 µg DNA. Cells were replated at different densities on gelatinised 9 cm dishes and subject to selection from 24-48 hours post-nucleofection. After 7-10 days, colonies were screened under a widefied microscope for red and/or green fluorescence, manually picked, dissociated with Trypsin-EDTA 0.05%, and replated into gelatinised 96-well plates. When confluent, cells were passaged to larger culture vessels, prior to expansion, banking and screening.

For lipofections, 2×10^5^ mESCs were plated onto gelatinised wells of a 6-well plate. Following full adhesion to the culture plate, they were lipofected with 2-5 µg DNA mixed with 2-5 µl/ug DNA P3000 solution and 2-5 µl Lipofectamine 3000 in OptiMEM.

### Cell lines

E14Ju09 ESCs are a 129/Ola male wild-type clonal line derived from E14tg2a (Hamilton and Brickman, 2014; Hooper et al., 1987).

Previously published SyNPL receiver (“STC”) and sender (“CmGP1”) cells are derived from E14Ju09 mESCs. STC cells contain randomly integrated *pPyCAG-tagBFP-3xNLS-IRES-Hph* and *pPGK-Signal-Myc-LaG17-mouse Notch1 core-tTA-IRES-Ble* constructs, and a *Rosa26*-targeted *TRE-mCherry* cassette. CmGP1 mESCs contain a randomly integrated *pPyCAG-EGFP-TMD-IRES-Pac* construct.

To generate KELPE cells, E14Ju09 cells were first nucleofected with linearised mKate2 landing pad construct to generate clonal mKate2-only landing pad cells. These cells were then nucleofected with linearised EGFP landing pad construct to generate clonal cell lines with two landing pads. To generate the final clonal KELPE cell line, the Vlox2272-SA-Hph-hGHpA-Vlox2272 resistance cassette present in the EGFP landing pad was excised by transiently transfecting a circular *pIns_CAG-VCre-P2A-HA-emiRFP670-3xNLS* plasmid, plating cells at clonal density, picking clones, replica plating them, and testing them for susceptibility to Hygromycin. It is also possible to sort single emiRFP670+ cells 24-48 hours after transfection to enrich for cells that have lost the Hygromycin resistance cassette. Correct targeting was confirmed by genomic DNA PCR with *OneTaq* polymerase, as shown in Tables S13-S17 (NB: if repeating these reactions, it is essential to use *OneTaq* polymerase. We have attempted these PCR reactions unsuccessfully with several other Taq polymerases) (Fig. S1A,B). KELPE cells were karyotyped by the IRR cell culture facility by counting chromosomes in metaphase spreads (Fig. S1C), then banked.

Untagged and tagged PUFFHalo cells were generated by lipofecting the KELPE cells with *CAG-phiC31 integrase* (Monetti et al., 2011) (a kind gift from Andras Nagy) and the appropriate RMCE construct: *AttB53-SA-Pac-SV40pA-cHS4(+)-PyF101-CAG-s36GFP-HaloTag-SpA-AttB53, AttB53-SA-Pac-SV40pA-cHS4(+)-PyF101-CAG-s36GFP-HaloTag-Flag-SpA-AttB53, AttB53-SA-Pac-SV40pA-cHS4(+)-PyF101-CAG-s36GFP-HaloTag-HA-SpA-AttB53, AttB53-SA-Pac-SV40pA-cHS4(+)-PyF101-CAG-s36GFP-HaloTag-Myc-SpA-AttB53* or *AttB53-SA-Pac-SV40pA-cHS4(+)-PyF101-CAG-s36GFP-HaloTag-V5-SpA-AttB53.*

KELPE-derived SyNPL receiver cells were generated by nucleofecting KELPE cells with *CAG-phiC31 integrase* and *AttB53-SA-Pac-bGHpA-cHS4(+)-PyF101-CAG-tagBFP-3xNLS-SpA-AttB53,* recovering puromycin-resistant tagBFP-positive mKate2-negative clones, and nucleofecting these with *pIns_CAG-FLPo-T2A-HA-emiRFP670* and *FRT5-Signal-Myc-LaG17-mouse Notch1 core-tTA-IRES-Bsd-SpA-cHS4(+)-(TRE-mCherry-SV40pA)(-)-FRT3*.

KELPE-drecived SyNPL sender cells were generated by nucleofecting KELPE cells with *pIns_CAG-FLPo-T2A-HA-emiRFP670* and the appropriate RMCE construct: *FRT5-EGFP-TMD-IRES-Bsd-SpA-FRT3* or *FRT5-EGFP-GPI-IRES-Bsd-SpA-FRT3*.

The inducible DTA receiver cells were generated by nucleofecting the KELPE cells with *CAG-phiC31 integrase* and *AttB53-SA-Pac-bGHpA-cHS4(+)-PyF101-CAG-tagBFP-3xNLS--SpA-AttB53,* recovering puromycin-resistant tagBFP-positive mKate2-negative clones, and nucleofecting these with *pIns_CAG-FLPo-T2A-HA-emiRFP670* and *FRT5-Signal-Myc-LaG17-mouse Notch1 core-tTA-IRES-Bsd-SpA-cHS4(+)-(TRE-DTA-SV40pA)(-)-FRT3*.

### Micropatterning

Micropatterning was performed as previously described (Robles-Garcia et al., 2024). Micropatterning was carried out in ibidi 8-well µSlides with an Alveole PRIMO bioengineering platform mounted on a Nikon Ti2 widefield microscope using a protocol adapted from Alveole. The surface of ibidi slides was first passivated as follows: slides were plasma treated (Harrick Plasma Cleaner) for 1 min 30 seconds at high intensity in a 200 mTorr vacuum and then incubated at room temperature for 1 h with 200 µl/well of 200 µg/ml poly-D-lysine diluted in 0.1 M HEPES pH 8.4. The wells were washed twice with ddH_2_O and once with 0.1 M HEPES pH 8.4 and then incubated for 1 h in the dark with 125 µl/well of 80 mg/ml mPEG-SVA freshly dissolved in 0.1 M HEPES pH 8.4. Slides were then washed profusely with ddH_2_O, air-dried and stored at 4°C until further processing (1 week maximum). The PRIMO insolation step was performed immediately prior to plating the cells: passivated wells were covered with 8 µl PLPP gel (1 µl PLPP gel/well diluted with 7 µl 70% ethanol) in the dark and left to dry for ∼30 min at room temperature. Slides were then insolated with PRIMO through a 20× lens with a dose of 50 mJ/cm^2^. The kelpie micropattern shape was designed in Adobe Illustrator and saved as a binary TIFF file prior to loading in the Alveole Leonardo software. After insolation, PLPP gel was removed with three ddH2O washes. Dulbecco’s phosphate buffered saline solution with calcium and magnesium (PBS++) was added to the wells for 5 minutes. Matrix coating was then performed by incubating the wells at room temperature for 30 min on a rocker with a mixture of 40 µg/ml vitronectin and 10 µg/ml laminin-521 diluted in PBS++. Wells were washed three times with PBS++ and left in the last wash whilst preparing the cells for seeding to ensure that the wells were not left to dry. For seeding, cells were passaged as normal and plated onto micropatterns at a density of 150,000 cells/well in 250 µl of ESC culture medium supplemented with 100 U/ml penicillin and 100 µg/ml streptomycin. The cells were left to adhere for 3 h at 37°C. After attachment, excess cells were removed by gentle pipetting and applying fresh ESC culture medium supplemented with penicillin/streptomycin. The cells were left to settle and cover the patterns overnight, before being live imaged the next day.

### Co-culture experiments

Cells were passaged as described in the “Mouse embryonic stem cell culture” section. For N2B27 experiments, cells were washed twice in N2B27 prior to final resuspension, to remove all traces of serum. Cells were counted manually or with a BioRad TC20 automated counter, and plated at the ratios described in the figure legends at a density of 2.5×10^5^ cells/cm^2^ onto cell culture vessels coated with 7.5 µg/ml fibronectin. Cells were co-cultured for 2 days prior to further analysis. For DTA induction experiments, we adapted the experimental timeframe to allow enough time for DTA to kill cells. It can take longer than 3 days for diphtheria toxin to kill mammalian cells (Zhang et al., 2010), so we carried out co-cultures over a period of 6 days to allow extra time for DTA transcription, translation and folding. We set up co-culture experiments as described above, and passaged cells on day 2 and day 4 (in order to maintain consistent density and facilitate contact between sender and receiver cells) prior to analysis on day 6.

### Flow cytometry

Cells were washed in PBS, detached from their culture vessel using Accutase, quenched in culture medium, pelleted by spinning at 300g for 3 min, and resuspended in ice cold PBS+10% FCS+live/dead stain. For PUFFHalo cells, 100 ng/ml DAPI was used, for KELPE and SyNPL cells, 300 nM DRAQ7 was used. Cells were filtered through a 30 µm cell strainer to reduce the number of doublets. Cells were analysed on a BD LSRFortessa flow cytometer. Single cells were identified using forward and side-scatter width and amplitude; dead cells were excluded by gating on either DAPI- or DRAQ7-negative cells. Then, fluorescence expression was analysed in live cells using the following laser/filter combinations: V 450/50-A (tagBFP), B 530/30-A (EGFP), Y/G 610/20-A (mCherry/mKate2), R 670/14-A (PUFFHalo-JF646). The data was analysed using the free online software Floreada (<u>https://floreada.io/</u>) in order to gate relevant populations, generate density plots and to export fluorescence data for gated samples (e.g. for “Live cells” or “Receiver cells”). The percentage of cells expressing each fluorophore was calculated by setting gates on negative controls. For experiments with PUFFHalo lines, adherent cultures were incubated in ESC culture medium supplemented with 10 nM Janelia Fluor 646 HaloTag Ligand (“JF646”) for 2 hours at 37°C and 5% CO2 prior to analysis.

To generate Violin SuperPlots, we first performed biexponential transformation of fluorescence data using the following formula:

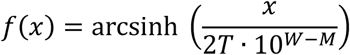

where T is the top of the scale, W is the width basis and M is the decade width. Values of each variable were empirically determined to avoid cropping of data. Values used were: T=262144, M=4.5, W=0.9 for all Violin SuperPlots with the exception of the mCherry plots, for which W=0.5.

This transformation, commonly used in commercial flow cytometry analysis software, is necessary to visualise negative fluorescence values generated by flow cytometers alongside a positive “log-like” region of the graph axes.

Violin SuperPlots were then generated in Python as described in the original publication (Kenny and Schoen, 2021).

### Immunofluorescence

Cells were plated on Ibidi 8 Well µ-Slides coated with 7.5 µg/ml fibronectin, and cultured as described in the figure legends. On the day of analysis, cells were washed with PBS, fixed in 4% formaldehyde in PBS for 20 min at room temperature. Cells were then subject to three 5-minute PBS washes. Cells were blocked either for 1 hr at room temperature or overnight at 4°C in blocking solution (PBS + 3% donkey serum + 0.1% Triton X-100). Primary antibodies were diluted in blocking solution and added for 3 hrs at room temperature or overnight at 4°C. After three 10-minute PBS washes for a total of minimum 30 min, secondary antibodies diluted in blocking solution were added for 1 hr at room temperature. Finally, cells were subject to three 10-minute PBS washes. PBS was replaced with a 90% glycerol aqueous solution for imaging and storage.

### Imaging

Live imaging of micropatterned KELPEs was performed on a widefield Nikon Ti-E microscope, with a 20X lens. Images were acquired with a Hamamatsu camera, using the Nikon NIS-Elements software to automate capture and stitching of an array of multiple fields of view.

Fixed and stained cells were imaged on a widefield Nikon Ti-E microscope, 20X lens and Hamamatsu camera, with the exception of the images in Fig. S2, which were acquired with a Leica SP8 confocal microscope with a 40X immersion lens.

### Statistical analysis

Definition of statistical significance, size of n, and statistical analysis performed is indicated in Figure Legends.

## ACKNOWLEDGEMENTS

We thank Sally Lowell, in whose lab some of this work was performed. We thank all users who have independently tested KELPE cell lines and their derivatives in our own laboratory as well as the laboratories of Sally Lowell, Leonardo Morsut, Marta Shahbazi and Elise Cachat; in particular Alicia Perez Lezcano, Catcher Salazar, Eleanor Earp, Eva Gonzalez Suarez, Jennifer Annoh, Hannah Coveney, Kiara Ramnundlall, Louise Chapman, Nanami Satoh, Peter Donlon, Maria Rosa Portero Migueles, Rupa Beke, Sophie Brumm, Tamina Lebek and Zunaira Aman. We also thank Tamina Lebek for help generating diagrams and for comments on the manuscript. We thank Sally Lowell and Sophie Brumm for comments on the manuscript. We thank Matthew French for assistance with micropatterning, Miguel Robles Garcia for micropatterning advice, and Guillaume Blin for micropatterning reagents. We thank Steven Pollard, Charles Williams and Rachel White for providing the YCe3736_HC_Amp_ccdB EMMA receiver vector and for advice on NSC differentiation. We thank Lucy Doyle for advice on NSC differentiation. We thank Andras Nagy, Anna-Katerina Hadjantonakis, James Briscoe and Wilson Wong for providing DNA constructs. We thank Matthieu Vermeren (IRR Imaging Facility), Justyna Cholewa-Waclaw (IRR High Content Screening Facility), David Kelly and Toni McHugh (Wellcome Discovery Research Platform for Hidden Cell Biology Light Microscopy Core) and Charles Williams for imaging support. We thank Fiona Rossi, Ailsa Laird, Claire Cryer, Aaron Hay, Anna Popravko and John Sutherland (IRR Flow Cytometry facility) for flow cytometry support. We thank Theresa O’Connor, Helen Henderson, Marilyn Thomson and Morag Heart for cell culture and karyotyping support.

## FOOTNOTES

## Author contributions

Conceptualization: A.F., M.M.; Methodology: A.F., M.M.; Validation: A.F., Y.S., M.M.; Formal analysis: A.F., Y.S., M.M.; Investigation: A.F., Y.S., M.M.; Resources: A.F., Y.S., M.M.; Data curation: A.F., Y.S., M.M.; Writing – Original Draft: A.F., M.M.; Writing – Review and Editing: A.F., Y.S., M.M.; Visualization: A.F., M.M.; Supervision: M.M.; Project administration: M.M.; Funding acquisition: M.M.

## Resource availability

Requests for resources described in this manuscript should be addressed to the corresponding author. Plasmids that were generated for this study will be made available on public repositories following peer-review and publication.

