## Supplementary Material for "KELPE: knock-in exchangeable dual landing pad embryonic stem cells enable efficient screening of synthetic gene circuits"

#### SUPPLEMENTARY INFORMATION

consisting of:

Table of contents

Supplementary Tables S1-S17

Supplementary Figures S1-S6

#### Table of contents

Table S1: Table S1: Site-specific recombinases and SSR sites

Table S2: Reagents

Table S3: Assembly of EGFP landing pad

Table S4: Assembly of untagged PUFFHalo RMCE vector

Table S5: Table S4: Assembly of tagged PUFFHalo RMCE vectors

Table S6: Assembly of tagBFP-3xNLS RMCE vector

Table S7: Assembly of synNotch-TRE-mCherry RMCE vector

Table S8: Assembly of synNotch-TRE-DTA RMCE vector

Table S9: Assembly of EGFP-TMD RMCE vector

Table S10: Assembly of EGFP-GPI RMCE vector

Table S11: Assembly of VCre recombinase expression vector

Table S12: Assembly of FLPo recombinase expression vector

Table S13: Primer sequences

Table S14: Genomic PCR Reaction contents – wild-type locus

Table S15: Genomic PCR Reaction contents – targeted locus (suitable for either landing pad)

Table S16: Genomic PCR Cycling conditions – wild-type locus

Table S17: Genomic PCR Cycling conditions – targeted locus (suitable for either landing pad)

Figure S1: KELPE cell validation

Figure S2: Maintenance of transgene expression following long-term culture and differentiation

Figure S3: EMMA assembly strategy

Figure S4: Validation of KELPE-derived PUFFHalo secretor cells

Figure S5: Validation of KELPE-derived SyNPL receiver cells

Figure S6: No DTA control co-culture experiment

#### Table S1: Site-specific recombinases and SSR sites

| Site-specific recombinase | SSR sites | Landing pad |
|---|---|---|
| phiC31 integrase | attP50 / attB53 | mKate2 landing pad |
| VCre | Vlox2272 / VloxP | EGFP landing pad |
| Dre | roxP / rox12 | EGFP landing pad |
| FLPo | FRT5 / FRT3 | EGFP landing pad |

#### Table S2: Reagents

| Reagent | Source | Identifier | Concentration |
|---|---|---|---|
| **Antibodies, nuclear counterstains, dyes** | | | |
| Rabbit polyclonal anti-HaloTag | Promega | Cat# G9281; RRID:AB_713650 | 1:200 |
| Mouse monoclonal anti-Flag | Sigma-Aldrich | Cat# F3165; RRID: AB_259529 | 1:1000 |
| Mouse monoclonal anti-HA | Sigma-Aldrich | Cat# H3663; RRID:AB_262051 | 1:1000 |
| Mouse monoclonal anti-Myc | Santa Cruz | Cat# sc-40; RRID:AB_627268 | 1:200 |
| Mouse monoclonal anti-V5 | Invitrogen | Cat# 14-6796-82; RRID:AB_10718239 | 1:200 |
| Chicken polyclonal anti-GFP | Abcam | Cat# ab13970; RRID:AB_300798 | 1:1000 |
| Rabbit polyclonal anti-tRFP (tagBFP, mKate2) | Evrogen | Cat# AB233; RRID:AB_2571743 | 1:1000 |
| Camelin sdAb anti-tagFP (tagBFP, mKate2), FluoTag x2 AF647 conjugated | NanoTag Biotechnologies | Cat# N0502-AF647-L; RRID:AB_3075936 | 1:200 |
| Rat monoclonal anti-mCherry | Invitrogen | Cat# M11217; RRID:AB_2536611 | 1:1000 |
| Mouse monoclonal anti-Oct3/4 | Santa Cruz | Cat# sc-5279; RRID:AB_628051 | 1:200 |
| Rat monoclonal anti-Krt8 | DSHB | Cat# TROMA-I; RRID:AB_531826 | 1:10 |
| Mouse monoclonal anti-Nestin | DSHB | Cat# Rat-401; RRID:AB_2235915 | 1:50 |
| DAPI | Biotium | Cat# BT40043 | 100 ng/ml (FACS) 1 µg/ml (IF) |
| DRAQ7 | Invitrogen | Cat# D15106 | 300 nM (FACS) 1 µM (IF) |
| Janelia Fluor 646 HaloTag Ligand | Promega | Cat# GA1121 | 10 nM |

| | | | |
|---|---|---|---|
| **Chemicals, solutions and competent bacteria** | | | |
| 1kb DNA ladder | NEB | Cat #N3232S | 600 ng/gel lane |
| 2-mercaptoethanol | Gibco | Cat# 31350010 | 100 nM |
| Accutase | Invitrogen | Cat# 00-4555-56 | |
| B-27 Supplement (50x) | Gibco | Cat# 17504-044 | 0.5X |
| Bovine Albumin Fraction V (7.5% solution) | Gibco | Cat# 15260-037 | 0.012% |
| Cultrex 3-D Culture Matrix Laminin I | Bio-Techne | Cat# 3446-005-01 | 2 μg/ml |
| D-(+)-Glucose solution | Sigma-Aldrich | Cat# G8644 | 8 mM |
| DMEM, no phenol red | Gibco | Cat# 31053028 | |
| DMEM/F12 | Gibco | Cat# 21331020 | |
| DMEM/HAMS-F12 | Sigma-Aldrich | Cat# D8437 | |
| Donkey serum | Sigma-Aldrich | Cat# D9663 | 3% v/v |
| Doxycycline hyclate | Sigma-Aldrich | Cat# D9891 | 1 μg/ml |
| Dulbecco's phosphate buffered saline (PBS) | Gibco | Cat# 14190-094 | |
| Dulbecco's phosphate buffered saline (PBS) with Calcium and Magnesium | Gibco | Cat# 1404141 | |
| Fibronectin from bovine plasma solution | Sigma-Aldrich | Cat# F1141 | 7.5 μg/ml |
| Foetal calf serum | Biosera | Cat# FB-1280/500 | 10% v/v |
| Formaldehyde 37-41% | Fisher Scientific | Cat# F/1501/PB08 | 4% v/v |
| Gelatin | Sigma Aldrich | Cat# G1890 | 0.1% w/v |
| Geneticin (G418) | Gibco | Cat# 11811031 | 200 μg/ml |
| Glasgow Minimum Essential Medium | Sigma Aldrich | Cat# G5154 | |
| Glycerol | Fisher Scientific | Cat# 15473719 | |
| HEPES sodium salt | Sigma Aldrich | Cat# H3784-25G | 0.1 M |
| Human FGF-basic | PeproTech | Cat# AF-100-18B | 10 ng/ml |
| Hygromycin B | Gibco | Cat# 10687010 | 200 μg/ml |
| L-Glutamine | Gibco | Cat# 25030024 | 2 mM |
| LIF | Made in-house | | 100 U |
| Lipofectamine 3000 transfection reagent | Invitrogen | Cat# L3000008 | |
| MEM non-essential amino acids solution (100x) | Gibco | Cat# 11140050 | 1X |
| Mouse EGF Recombinant protein | PeproTech | Cat# 315-09 | 10 ng/ml |
| mPEG-SVA | Laysan Bio | Cat# MPEG-SVA-5000 | 80 mg/ml |
| N-2 Supplement (100x) | Gibco | Cat# 17502-048 | 0.5X |
| NEB® 5-alpha Competent *E. coli* (Subcloning Efficiency) | NEB | Cat# C2988J | |

| | | | |
|---|---|---|---|
| NEB® Stable Competent E. coli (High Efficiency) | NEB | Cat# C3040I | |
| NEBridge® Golden Gate Assembly Kit (BsmBI-v2) | NEB | Cat# E1602S | |
| Neurobasal medium | Gibco | Cat# 21103049 | |
| One*Taq*® Hot Start DNA Polymerase | NEB | Cat# M0481S | |
| Penicillin/streptomycin | Gibco | Cat# 15140122 | 1:100 (100 U/ml penicillin, 100 µg/ml streptomycin) |
| PLPP gel | Alveole | Cat# PLPP_Gel | 1:8 |
| Poly-D-Lysine Hydrobromide | Merck Millipore | Cat# A-003-E | 200 µg/ml |
| Puromycin dihydrochloride | Sigma Aldrich | Cat# P8833 | 2 µg/ml |
| PD0325901 | Axon Medchem | Cat# 1408 | 0.5 µM |
| Recombinant human Laminin-521 | Gibco | Cat# A29249 | 10 µg/ml |
| Sodium pyruvate | Gibco | Cat# 11360039 | 1 mM |
| Triton X-100 | Sigma Aldrich | T8787 | 0.1% v/v |
| Trypsin-EDTA 0.05% | Gibco | Cat# 25300054 | |
| Vitronectin (VTN-N) Recombinant Human Protein, Truncated | Thermo Fisher | Cat# A14700 | 40 µg/ml |
| Zeocin | Gibco | Cat# R25001 | 100 µg/ml |
| Zero Blunt™ TOPO™ PCR Cloning Kit, without competent cells | Invitrogen | Cat# 450245 | |
| **Cell culture vessels** | | | |
| µ-Slide 8 Well | Ibidi | Cat# 80826 | |
| Plastic flasks and plates for routine cell culture | Corning | Various | |
| **DNA constructs** | | | |
| pmROSA26-attP50-Neo-mKate2-3xNLS-attP50 plasmid (Addgene plasmid #183609) | Malaguti et al., 2022 | Addgene 183609 | |
| pmROSA26-ANCKAC (mKate2 landing pad) | This study | | |
| pmROSA26-HELP (EGFP landing pad) | This study | | |
| GBSGFP insulator | Balaskas et al., 2012 | Addgene 121975 | |

| | | |
|---|---|---|
| CAG-phiC31 integrase | Monetti et al., 2011 | Similar plasmid available: Addgene 62658 |
| pCAG-VCre | Weinberg et al., 2017 | Addgene 89575 |
| pIns_CAG-VCre-P2A-HA-emiRFP670-3xNLS | This study | |
| pCAG-FlpO | Weinberg et al., 2017 | Addgene 89574 |
| pIns_CAG-FLPo-P2A-HA-emiRFP670-3xNLS | This study | |
| APCCBA (AttB53-SA-Pac-bGHpA-cHS4-PyCAG-tagBFP-3xNLS-SpA-AttB53) | This study | |
| AttB53-SA-Pac-bGHpA-cHS4-PyCAG-PUFFHalo-SpA-AttB53 | This study | |
| AttB53-SA-Pac-bGHpA-cHS4-PyCAG-PUFFHalo-Flag-SpA-AttB53 | This study | |
| AttB53-SA-Pac-bGHpA-cHS4-PyCAG-PUFFHalo-HA-SpA-AttB53 | This study | |
| AttB53-SA-Pac-bGHpA-cHS4-PyCAG-PUFFHalo-Myc-SpA-AttB53 | This study | |
| AttB53-SA-Pac-bGHpA-cHS4-PyCAG-PUFFHalo-V5-SpA-AttB53 | This study | |
| FRT5-Signal-Myc-LaG17-mN1C-tTA-IRES-Bsd-SpA-cHS4-(TRE-mCherry-SV40pA)(-)-FRT3 | This study | |
| FRT5-Signal-Myc-LaG17-mN1C-tTA-IRES-Bsd-SpA-cHS4-(TRE-DTA-SV40pA)(-)-FRT3 | This study | |
| FRT5-Signal-EGFP-TMD-IRES-Bsd-SpA-FRT3 | This study | |
| FRT5-Signal-EGFP-GPI-IRES-Bsd-SpA-FRT3 | This study | |
| pCAG:GPI-GFP | Rhee et al., 2006 | Addgene 32601 |
| **Cell lines** | | |
| E14Ju09 mouse ESCs (129/Ola, male) | Hamilton and Brickman, 2014 | |

| | |
|---|---|
| KELPE mouse ESCs (129/Ola, male) | This study |
| RRK mouse ESCs (mKate2 landing pad only) | This study |
| ELPR mouse ESCs (EGFP landing pad only) (129/Ola, male) | This study |
| GPH (KELPE-derived PUFFHalo mouse ESCs) (129/Ola, male) | This study |
| GPH-F (KELPE-derived PUFFHalo-Flag mouse ESCs) (129/Ola, male) | This study |
| GPH-H (KELPE-derived PUFFHalo-HA mouse ESCs) (129/Ola, male) | This study |
| GPH-M (KELPE-derived PUFFHalo-Myc mouse ESCs) (129/Ola, male) | This study |
| GPH-V (KELPE-derived PUFFHalo-V5 mouse ESCs) (129/Ola, male) | This study |
| CmGP1 (SyNPL sender mouse ESCs) (129/Ola, male) | Malaguti et al., 2022 |
| STC (SyNPL receiver mouse ESCs) (129/Ola, male) | Malaguti et al., 2022 |
| BSNIB (KELPE-derived SyNPL receiver inducible mCherry mouse ESCs) (129/Ola, male) | This study |
| BSNIB-DTA (KELPE-derived SyNPL receiver inducible DTA mouse ESCs) (129/Ola, male) | This study |
| EEtm (KELPE-derived SyNPL sender EGFP-TMD mouse ESCs) (129/Ola, male) | This study |
| EEgpi (KELPE-derived SyNPL sender EGFP-GPI mouse ESCs) (129/Ola, male) | This study |

#### Table S3: Assembly of EGFP landing pad

| EMMA Position | Part description |
|---|---|
| 1 | Vlox2272-SA-P2A-Hph-hGHpA-Vlox2272 |
| 2 | cHS4(+) |
| 3 | roxP |
| 4 | CAG promoter |
| 5 | FRT5 |
| 6-8 | EGFP-3xNLS |
| 9-20 | IRES-Ble-rbGpA |
| 21 | FRT3 |
| 22 | rox12 |
| 23 | cHS4(-) |
| 24 | VloxP |
| 25 | AscI-FspAI restriction sites |
| Backbone | EMMA receiver vector |

#### Table S4: Assembly of untagged PUFFHalo RMCE vector

| EMMA Position | Part description |
|---|---|
| 1 | attB53 |
| 2 | SA-Pac-bGHpA-cHS4(+) |
| 3 | PyF101 enhancer |
| 4 | CAG promoter |
| 5-10 | PUFFHalo |
| 11-22 | Synthetic polyA |
| 23-24 | Assembly connector |
| 25 | attB53 |
| Backbone | EMMA receiver vector |

#### Table S5: Assembly of tagged PUFFHalo RMCE vectors

| EMMA Position | Part description |
|---|---|
| 1 | attB53 |
| 2 | SA-Pac-bGHpA-cHS4(+) |
| 3 | PyF101 enhancer |
| 4 | CAG promoter |
| 5-10 | PUFFHalo |
| 11-12 | Protein tag |
| 13-22 | Synthetic polyA |
| 23-24 | Assembly connector |
| 25 | attB53 |
| Backbone | EMMA receiver vector |

#### Table S6: Assembly of tagBFP-3xNLS RMCE vector

| EMMA Position | Part description |
|---|---|
| 1 | attB53 |
| 2 | SA-Pac-bGHpA-cHS4(+) |
| 3 | PyF101 enhancer |
| 4 | CAG |
| 5-10 | tagBFP-3xNLS |
| 11-22 | Synthetic polyA |
| 23-24 | Assembly connector |
| 25 | attB53 |
| Backbone | EMMA receiver vector |

#### Table S7: Assembly of synNotch-TRE-mCherry RMCE vector

| EMMA Position | Part description |
|---|---|
| 1-4 | Assembly connector |
| 5-6 | FRT5 |
| 7-10 | synNotch receptor (LaG17-mNotch1 core-tTA) |
| 11 | IRES |
| 12 | Bsd |
| 13-22 | Synthetic polyA |
| 23 | cHS4(+) |
| 24 | TRE-mCherry-SV40pA (-strand) |
| 25 | FRT3 |
| Backbone | EMMA receiver vector |

#### Table S8: Assembly of synNotch-TRE-DTA RMCE vector

| EMMA Position | Part description |
|---|---|
| 1-4 | Assembly connector |
| 5-6 | FRT5 |
| 7-10 | synNotch receptor (LaG17-mNotch1 core-tTA) |
| 11 | IRES |
| 12 | Bsd |
| 13-22 | Synthetic polyA |
| 23 | cHS4(+) |
| 24 | TRE-DTA-SV40pA (-strand) |
| 25 | FRT3 |
| Backbone | EMMA receiver vector |

#### Table S9: Assembly of EGFP-TMD RMCE vector

| EMMA Position | Part description |
|---|---|
| 1-4 | Assembly connector |
| 5-6 | FRT5 |
| 7-10 | Mouse IgGk signal sequence – EGFP – human PDGFB transmembrane domain |
| 11 | IRES |
| 12 | Bsd |
| 13-22 | Synthetic polyA |
| 23 | cHS4(+) |
| 24 | TRE-DTA-SV40pA (-strand) |
| 25 | FRT3 |
| Backbone | EMMA receiver vector |

#### Table S10: Assembly of EGFP-GPI RMCE vector

| EMMA Position | Part description |
|---|---|
| 1-4 | Assembly connector |
| 5-6 | FRT5 |
| 7-10 | Mouse acrosin signal sequence – EGFP – mouse Thy1 GPI signal sequence |
| 11 | IRES |
| 12 | Bsd |
| 13-22 | Synthetic polyA |
| 23 | cHS4(+) |
| 24 | TRE-DTA-SV40pA (-strand) |
| 25 | FRT3 |
| Backbone | EMMA receiver vector |

#### Table S11: Assembly of VCre recombinase expression vector

| EMMA Position | Part description |
|---|---|
| 1 | Assembly connector |
| 2 | cHS4(+) |
| 3 | PyF101 enhancer |
| 4 | CAG promoter |
| 5-12 | VCre |
| 13 | P2A |
| 14 | HA-emiRFP670-3xNLS |
| 15 | Synthetic polyA |
| 16-25 | cHS4(+) |
| Backbone | EMMA receiver vector |

#### Table S12: Assembly of FLPo recombinase expression vector

| EMMA Position | Part description |
|---|---|
| 1 | Assembly connector |
| 2 | cHS4(+) |
| 3 | PyF101 enhancer |
| 4 | CAG promoter |
| 5-12 | FLPo |
| 13 | P2A |
| 14 | HA-emiRFP670-3xNLS |
| 15 | Synthetic polyA |
| 16-25 | cHS4(+) |
| Backbone | EMMA receiver vector |

#### Table S13: Primer sequences

| Target | Primer sequence |
|---|---|
| Rosa26 wild-type | Forward : CAAAGTCGCTCTGAGTTGTTATCAG<br>Reverse : GGAGCGGGAGAAATGGATATGAAG |
| Rosa26 mKate2 landing pad | Forward : GAGCGGTGCAATGGTGTGTA<br>Reverse: CGATCCGTCGCCAAGCTTTTA |
| Rosa26 EGFP landing pad | Forward: GAGCGGTGCAATGGTGTGTA<br>Reverse: TAGCACGCGTGACGGTCG |

#### Table S14: Genomic PCR Reaction contents – wild-type locus

| Component | Volume | Concentration |
|---|---|---|
| 5X One*Taq* **Standard** Reaction Buffer | 5 µl | 1X |
| 10 mM dNTPs | 0.5 µl | 200 µM each |
| 10 µM Primer Forward | 0.5 µl | 200 nM |
| 10 µM Primer Reverse | 0.5 µl | 200 nM |
| *OneTaq* DNA polymerase | 0.25 µl | 1.25 units |
| Template genomic DNA | | 100 ng |
| Nuclease-free water | To 25 µl | |

#### Table S15: Genomic PCR Reaction contents – targeted locus (suitable for either landing pad)

| **Component** | **Volume** | **Concentration** |
|---|---|---|
| 5X One*Taq* **GC** Reaction Buffer | 5 µl | 1X |
| 10 mM dNTPs | 0.5 µl | 200 µM each |
| 10 µM Primer Forward | 0.5 µl | 200 nM |
| 10 µM Primer Reverse | 0.5 µl | 200 nM |
| *OneTaq* DNA polymerase | 0.25 µl | 1.25 units |
| Template genomic DNA | | 100 ng |
| Nuclease-free water | To 25 µl | |

#### Table S16: Genomic PCR Cycling conditions – wild-type locus

| **Step** | **Temperature** | **Time** |
|---|---|---|
| Initial denaturation | 94°C | 5 minutes |
| 35 cycles | 94°C | 30 seconds |
| | 58°C | 30 seconds |
| | 68°C | 1 minute |
| Final extension | 68°C | 5 minutes |
| Hold | 4-10°C | |

#### Table S17: Genomic PCR Cycling conditions – targeted locus (suitable for either landing pad)

| **Step** | **Temperature** | **Time** |
|---|---|---|
| Initial denaturation | 94°C | 5 minutes |
| 35 cycles | 94°C | 30 seconds |
| | 58°C | 30 seconds |
| | 68°C | 2 minutes |
| Final extension | 68°C | 5 minutes |
| Hold | 4-10°C | |

### FIGURE S1

#### A - Genomic PCR strategy

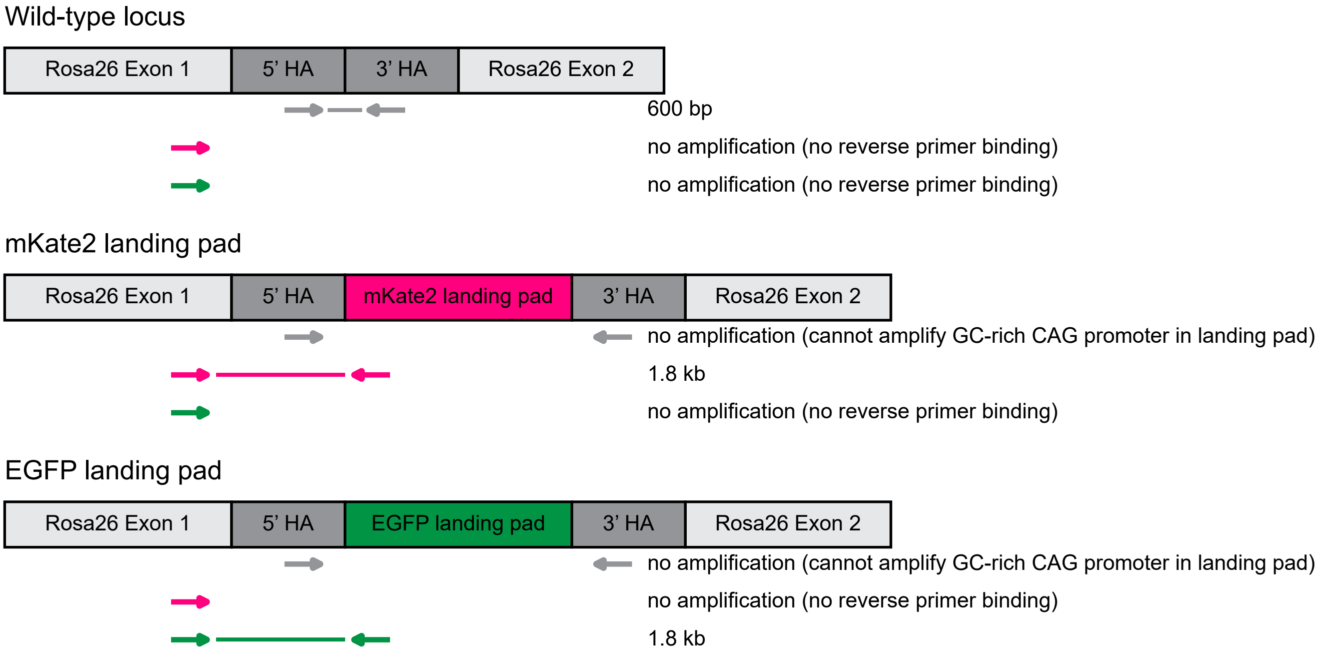

#### B - Genomic PCR result

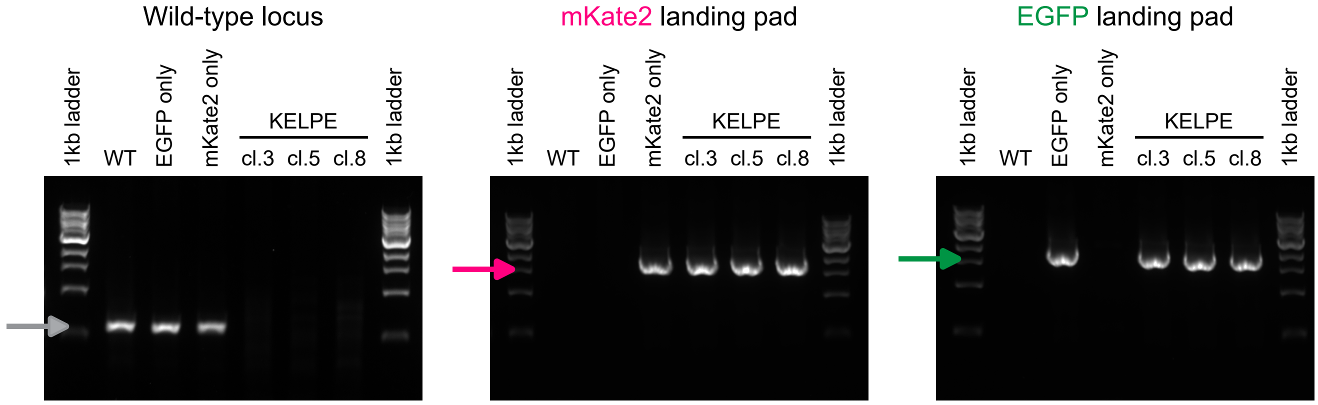

#### C - Karyotype

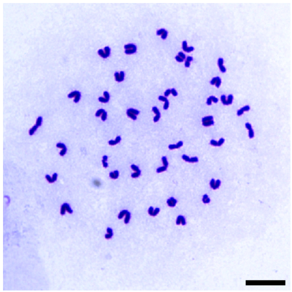

**Figure S1 - KELPE cell validation**
(A) Genomic PCR strategy to test for presence of the wild-type *Rosa26* locus, for correct integration of the mKate2 landing pad, and for correct integration of the EGFP landing pad. Expected band sizes are shown.
(B) Genomic PCR results for the PCRs shown in (A) for wild-type ("WT") cells, cells targeted with the EGFP landing pad but no mKate2 landing pad ("EGFP only"), cells targeted with the mKate2 landing pad but no EGFP landing pad ("mKate2 only"), and three KELPE clones. Samples were run alongisde the NEB 1kb ladder. Expected band sizes are shown as arrows.
(C) Representative metaphase spread of KELPE cl.5. Scale bar: 10 µm.

#### FIGURE S2

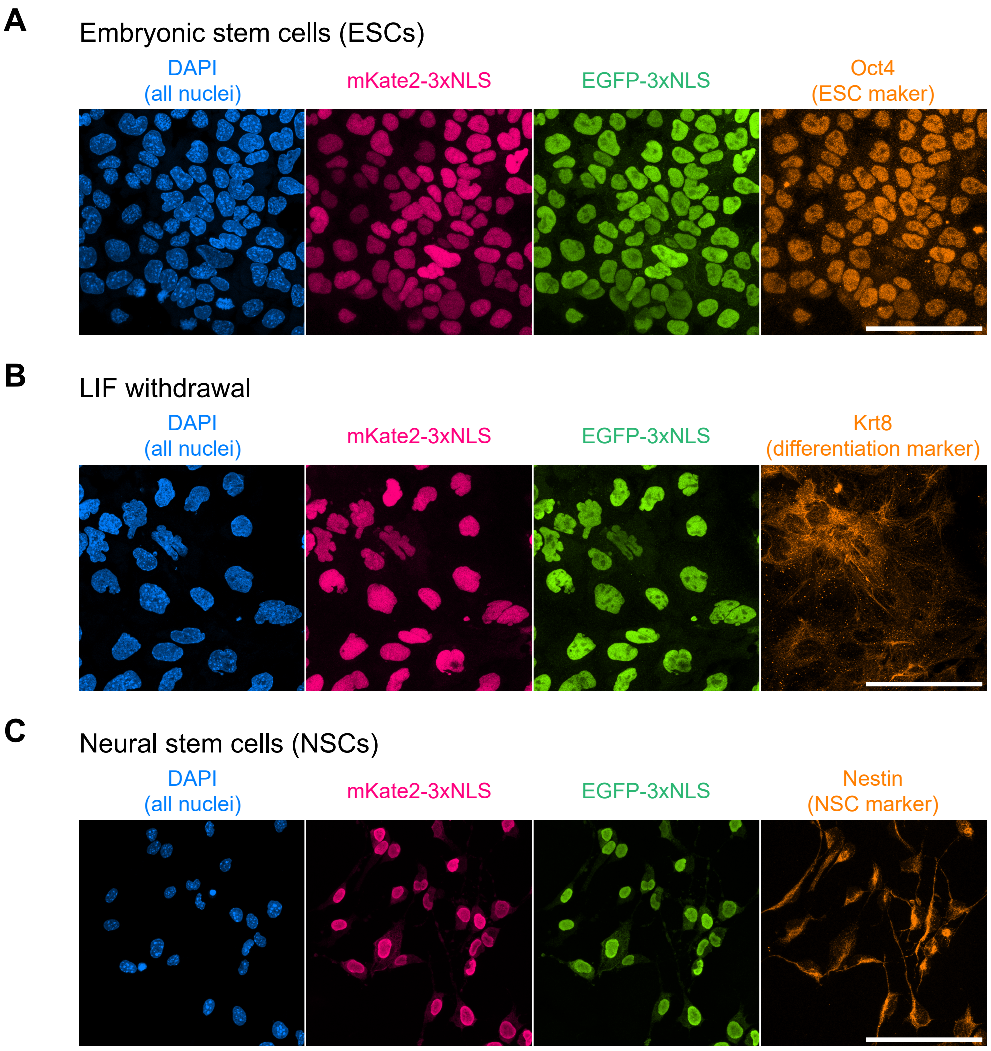

**Figure S2 - Maintenance of transgene expression following long-term culture and differentiation**
Immunofluorescence of KELPE cells cultured for >28 days in the absence of antibiotic selection.
(A) KELPE cells in ESC culture medium, stained for the pluripotency marker Oct4.
(B) KELPE cells in LIF withdrawal medium (multilineage differentiation), stained for the differentiation marker Krt8.
(C) KELPE cells differentiated into neural stem cells (NSCs), stained for the NSC marker Nestin.
Each panel was acquired separately, fluorescence intensities are not directly comparable. Scale bars: 100 μm.

#### FIGURE S3

#### EMMA assembly strategy

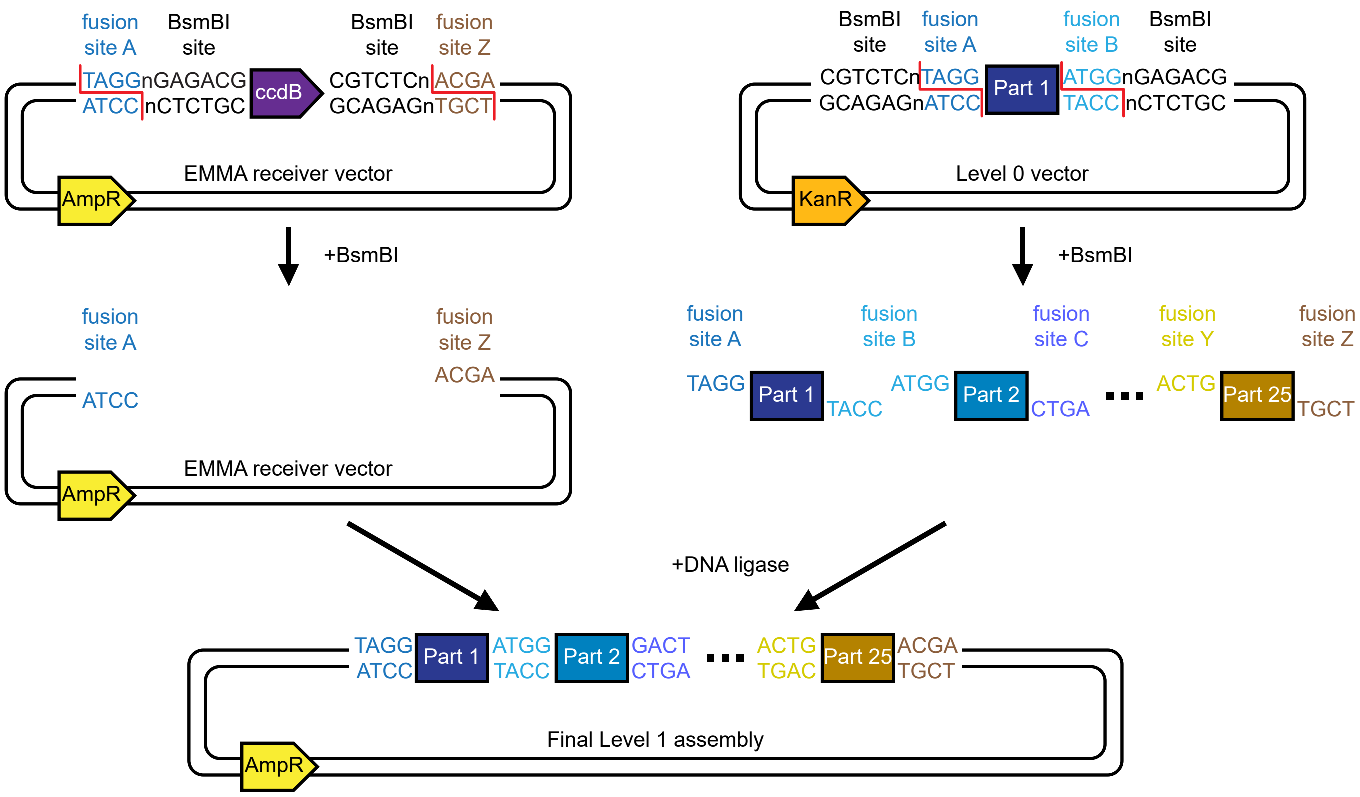

**Figure S3 - EMMA assembly strategy**
The EMMA receiver vector carries an ampicillin resistance gene (AmpR), as well as a toxic gene (*ccdB*) flanked by BsmBI sites. Digestion of the vector with BsmBI excises the *ccdB* toxic gene exposing two fusion sites (A and Z). Level 0 vectors carry a kanamycin resistance gene (KanR) and two BsmBI sites, which are placed in reverse orientation to the receiver vector, so that fusion sites are adjacent to a domesticated DNA part (e.g. Part 1). Digestion of Level 0 vectors with BsmBI releases the domesticated DNA part with its fusion site sequences. Ligation of the BsmBI-digested EMMA receiver vector with the BsmBI-digested DNA parts leads to assembly of the final Level 1 target construct, which confers ampicillin resistance and does not contain the *ccdB* toxic gene.

#### FIGURE S4

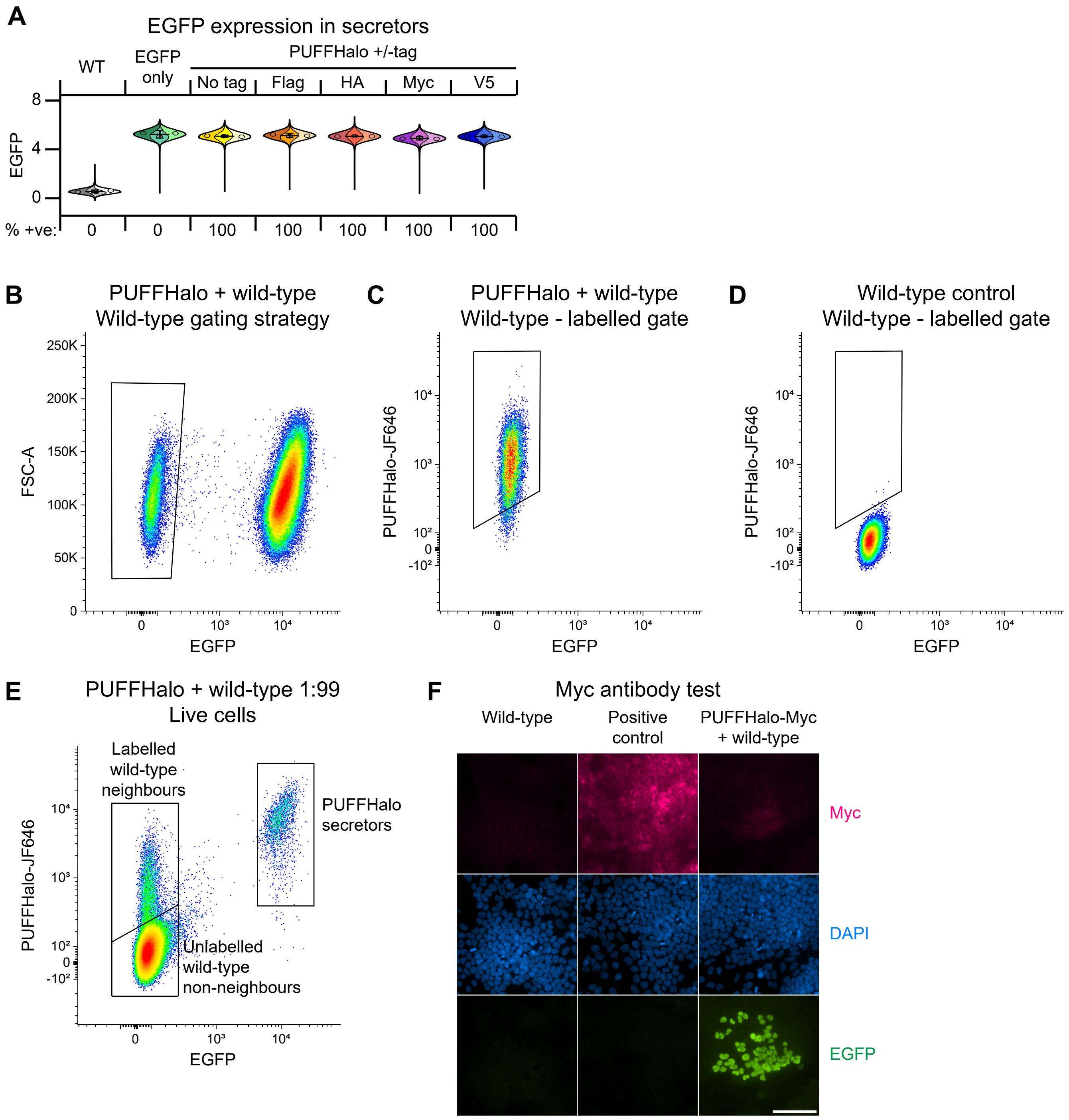

**Figure S4 - Validation of KELPE-derived PUFFHalo secretor cells**
(A) Flow cytometry analysis of EGFP expression in wild-type ("WT") cells, cells containing the EGFP-3xNLS landing pad but no mKate2 landing pad ("EGFP only"), and KELPE-derived PUFFHalo mESC lines. Data is displayed for 3 replicates (WT, EGFP only) or 3 clones (each PUFFHalo lines). The percentage of EGFP-positive cells was calculated using the gate shown in (B).
(B) Gating strategy to identify wild-type cells in co-culture with EGFP-positive PUFFHalo secretor cells.
(C) Gating strategy to identify labelled wild-type cells following co-culture with PUFFHalo secretor cells.
(D) Control sample showing wild-type cells cultured on their own, and the "labelled" gate shown in (C).
(E) Flow cytometry analysis of PUFFHalo secretor cells co-cultured with wild-type cells at 1:99 ratio. Gats are shown to illustrate the location of the plot of PUFFHalo secretors, labelled wild-type neighbours, and unlabelled wild-type neighbours.
(F) Anti-Myc antibody test immunofluorescence. Wild-type cells were used as a negative control, KELPE-derived SyNPL receiver cells, expressing Myc-tagged synNotch receptor, were used as a positive control, a 1:99 co-culture of PUFFHalo-Myc secretors with wild-type cells was used as a test sample. Scale bar: 100 µm.
A minimum of 9000 cells are displayed in each flow cytometry plot.

#### FIGURE S5

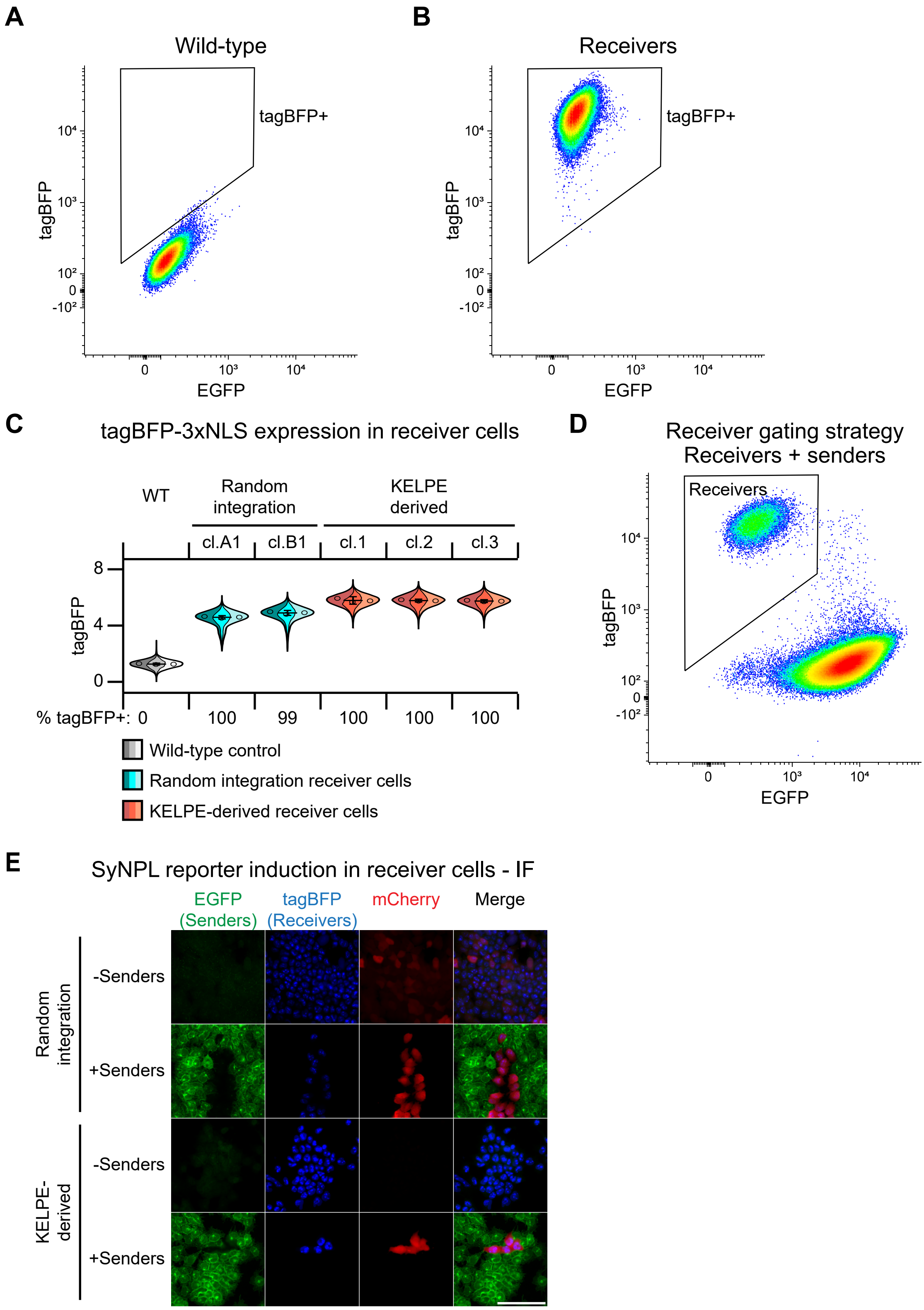

**Figure S5 - Validation of KELPE-derived SyNPL receiver cells**
(A) Flow cytometry analysis of EGFP and tagBFP expression in wild-type cells (negative control). These cells were used to set a "tagBFP+" gate to identify SyNPL receiver cells. (B) Flow cytometry analysis of EGFP and tagBFP expression in SyNPL receiver cells, which fall within the tagBFP+ gate. (C) Flow cytometry analysis of tagBFP expression in wild-type ("WT") cells, random integration SyNPL receiver cells, and KELPE-derived SyNPL receiver cells. 3 biological replicates are displayed for each cell line. Percentages of tagBFP-positive cells were calculated using the "tagBFP+" gate in (A) and (B). (D) Gating strategy to separate tagBFP+ receiver cells from EGFP+ sender cells by flow cytometry in co-culture experiments. (E) Culture of random integration SyNPL receiver cells (cl. A1) or KELPE-derived receiver cells (cl. 1) alone or in co-culture with sender cells (19:1 sender: receiver ratio). Immunofluorescence for EGFP, tagBFP and mCherry. Scale bar: 100 μm. A minimum of 10,000 cells are displayed in each flow cytometry plot.

#### FIGURE S6

Control (no DTA) receiver cells

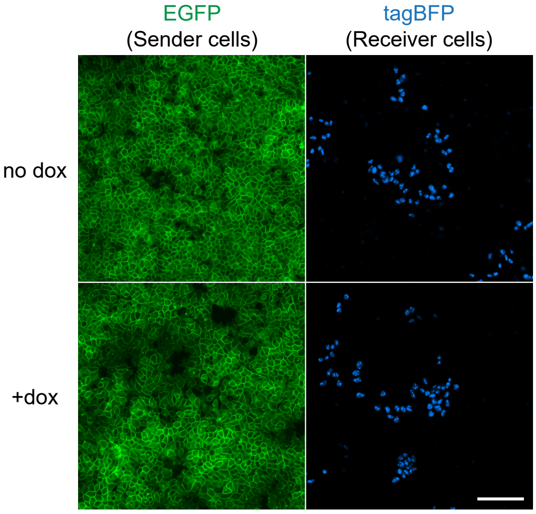

**Figure S6 - No DTA control co-culture experiment**
Immunofluorescence for EGFP and tagBFP of co-cultures of EGFP-expressing SyNPL sender cells with tagBFP-expressing control receiver cells that do not contain an inducible DTA transgene (19:1 sender:receiver ratio), in the absence (”no dox”) or presence (”+dox”) of doxycyline. Scale bar: 100 μm.
